# A chloroplast sulfate transporter modulates glutathione-mediated redox cycling to regulate cell division

**DOI:** 10.1101/2023.12.07.570675

**Authors:** Pin-jui Huang, Chun-Han Chen, Yen-Ling Lin, Hsiang-Yin Lin, Su-Chiung Fang

## Abstract

Glutathione redox cycling is important for cell cycle regulation. However, the underlying mechanisms remain unclear. We previously identified a cell-size mutant, *suppressor of mat3 15-1* (*smt15-1*), that has elevated cellular glutathione, increased number of cell divisions, and small daughter cells. Here, we demonstrated that SMT15 is a chloroplast-associated membrane protein that is capable of transporting sulfate. Reducing expression of *γ-GLUTAMYLCYSTEINE SYNTHETASE*, which encodes the rate-limiting enzyme required for glutathione biosynthesis, corrected the size defect of *smt15-1* cells. Moreover, overexpressing *GLUTATHIONE SYNTHETASE* recapitulated the small-size phenotype of *smt15-1* mutant, confirming the role of glutathione in modulation of the cell division. Hence, SMT15 may regulate chloroplast sulfate concentration to modulate cellular glutathione levels. Interestingly, glutathione was found to accumulate in the cytosol at the G1 phase and its level decreased substantially as cells entered the S/M phase in wild-type cells. Even though cytosolic glutathione of the small-sized mutants, *smt15-1* and *GSH2* overexpressors, followed the pattern of wild-type cells being accumulated at G1 and declined at the S/M phase, the basal body-specific accumulation of glutathione was associated with only the small-sized mutants. Therefore, we propose that glutathione-mediated redox in the basal bodies may regulate mitotic division number in *Chlamydomonas reinhardtii*. Our results support the link between glutathione-mediated redox regulation and mitotic cell division and suggest a new mechanism through which glutathione regulates the cell cycle.

**One sentence summary:** Glutathione-mediated redox regulation in basal bodies is important for cell division control

## Introduction

Reactive oxygen species (ROS) signaling plays an important role in cell cycle regulation in eukaryotic organisms (Burch and Heintz, 2005; Burhans and Heintz, 2009; Tsukagoshi et al., 2010; Chiu and Dawes, 2012; Dolzblasz et al., 2016; Yu et al., 2016; de Simone et al., 2017; Ivanova et al., 2021). Metabolic fluctuation between the oxidative and reductive phases is regulated as a function of the cell cycle with cell division strictly confined to the reductive phase in budding yeast (Tu et al., 2005). Having DNA replication restricted to the reductive phase of the metabolic cycle is to ensure genome integrity in budding yeast (Chen et al., 2007). The coupling of the cell cycle with redox-associated metabolic oscillation is also reported in the mammalian system (Sarsour et al., 2009; Vander Heiden et al., 2009; Chiu and Dawes, 2012; Ahn et al., 2017). In mammals, low levels of ROS are capable of activating cell cycle regulators and stimulating cell cycle entry (Lee et al., 1998; Martindale and Holbrook, 2002; Boonstra and Post, 2004; Macleod, 2008; Burhans and Heintz, 2009; Lyublinskaya et al., 2015; Kirova et al., 2022). Conversely, cell cycle regulators are also important to regulate redox-associated cell metabolism by directly regulating the pentose phosphate and serine pathways (Wang et al., 2017). Even though redox regulation of the cell cycle has become an emerging principle, knowledge of the underlying molecular mechanisms connecting the cell cycle to redox signaling remain limited.

Glutathione is a thiol-containing tripeptide that functions to maintain redox homeostasis in response to biotic and abiotic stresses in plants and animals (Cobbett et al., 1998; Ball et al., 2004; Wu et al., 2004; Mhamdi et al., 2010; Shanmugam et al., 2012; Shanmugam et al., 2015; Chen et al., 2016; Trujillo-Hernandez et al., 2020). Defects in glutathione-dependent redox balance also affect the cell cycle (Vernoux et al., 2000; Cairns et al., 2006; Odom et al., 2009; Jiao et al., 2013), suggesting that glutathione has a regulatory role in the cell cycle. Consistent with this notion, temporal and spatial regulation of glutathione is tightly associated with the cell cycle (Diaz Vivancos et al., 2010; Fang et al., 2014; de Simone et al., 2017; Hartl et al., 2020). Moreover, recent studies have revealed a mechanistic link between the negative cell cycle regulator - retinoblastoma tumor suppressor (RB) and alternated glutathione homeostasis (Nicolay et al., 2013; Mandigo et al., 2021). Despite these reports, how the glutathione-dependent redox state modulates cell division cycle remains poorly understood.

*Chlamydomonas reinhardtii*, herein referred to as Chlamydomonas, is a unicellular green alga that reproduces by a modified cell cycle named multiple fission. Multiple fission starts with a long G1 phase, during which cells can grow photosynthetically and reach sizes that can be several times that of the newborn (daughter) cells. At the end of G1, mother cells undergo n very quick rounds of alternating S phase (DNA synthesis) and M phase (mitosis) to produce 2^n^ daughter cells (Pecani et al., 2022). During multiple fission, two key control checkpoints are proposed to explain size homeostasis of the newborn daughters (Umen, 2005; Fang and Umen, 2008). In early/mid G1, cells pass the first size checkpoint, commitment, which requires cells to acquire a minimum size threshold for cells to complete the cell cycle. During S/M phase, the second size checkpoint, mother cells undergo a regulated number of division cycles regardless of their size to produce uniformly sized daughters (Craigie and Cavaliersmith, 1982; Donnan and John, 1983). Because *C. reinhardtii* mother cell division can occur in the absence of concurrent growth, daughter cell-size can be conveniently used to assess the cell size checkpoint function during S/M phase (Umen, 2005).

Previous studies have demonstrated that the Chlamydomonas Retinoblastoma (RB) pathway is important for cell-size checkpoint functions of commitment and S/M phase (Umen and Goodenough, 2001; Fang et al., 2006). The Chlamydomonas RB homolog is encoded by a single gene, *MAT3*. The *mat3* mutant cells pass commitment at a reduced cell-size and undergo an increased number of cell divisions to produce tiny daughter cells. In a cell-size suppression genetic screen, mutants that have defects in the Chlamydomonas *E2F1* or *DP1* genes (the direct downstream targets of animal RB protein) bypass *mat3* size defect and suppress the small-sized *mat3* to generate large cells, demonstrating that the architecture of Chlamydomonas RB pathway is evolutionarily conserved (Fang et al., 2006; Olson et al., 2010).

In addition to the canonical RB pathway, genetic screen isolated additional extragenic suppressors of *mat3* (*smt*) mutant whose size-suppression phenotype is weaker than that of *e2f1* and *dp1* mutants. (Fang and Umen, 2008; Lin et al., 2020). Among these *smt* suppressors, *smt15-1 mat3-4* double mutants are larger than *mat3-4* single mutant but smaller than wild-type cells, suggesting *SMT15* may be a positive regulator of cell division. Paradoxically, *smt15-1* single mutant has daughter cells that are smaller than those of the wild-type, indicating that SMT15 has a negative regulatory role in controlling size-dependent cell division. Together, the accumulated results suggest that SMT15 modulates the Chlamydomonas RB pathway through a novel mechanism. SMT15 belongs to a large family of putative sulfate/anion transporters that contain a sulfate transporter domain (Fang et al., 2014). In addition to the small size, *smt15-1* mutant also has elevated cellular glutathione content. In Chlamydomonas, glutathione level fluctuates during the cell cycle with highest level in mid G1 phase and lower level during S/M phases. Even though glutathione level is comparable in wild-type and *smt15-1* strains at G1 phase, glutathione level of *smt15-1* cells failed to decline as cells enter S/M phase (Fang et al., 2014). Together, these results suggest that glutathione may have a role in size-mediated cell cycle control and SMT15 regulates size-dependent cell division through modulating glutathione homeostasis.

In this study, we demonstrate that SMT15 is a chloroplast-associated membrane protein capable of transporting sulfate. Decreasing glutathione by reducing expression of the *γ-GLUTAMYLCYSTEINE SYNTHETASE* (*GSH1*), the rate-limiting enzyme for glutathione biosynthesis, restored the size defect of *smt15-1* mutant by increasing its daughter cell size, confirming that glutathione modulates the cell-size control mechanism of *smt15-1* mutant. Furthermore, altering glutathione homeostasis by overexpressing *GLUTATHIONE SYNTHETASE* (*GSH2*) recapitulated the small-size phenotype of *smt15-1* mutant, arguing again that glutathione regulates size-mediated cell cycle control. During the cell cycle, we found that glutathione was accumulated primarily in the cytosol at G1 phase and the cytosolic glutathione decreased substantially as cells enter S/M phase. Intriguingly, glutathione was found to be specifically concentrated in the basal bodies of *smt15-1* and *GSH2* overexpressors cells at the S/M phase. Hence, we suggested that glutathione redox regulation in the basal body may be a novel mechanism to regulate size-mediated cell division control in *C. reinhardtii*. Taking these results together, we propose a model to explain how a sulfate transporter SMT15 affects glutathione homeostasis and regulates cell-size checkpoint function.

## Results

### SMT15 encodes a sulfate transporter

Our previous data showed that *SMT15* encodes a potential sulfate transporter that belongs to tribe 1 of the eukaryotic sulfate/anion transporters (Fang et al., 2014). To investigate the sulfate transporter activity of SMT15, we attempted to amplify the full-length SMT15 cDNA for sulfate uptake assay. Unfortunately, the full-length SMT15 cDNA contains many GC repeats and is extremely GC rich (71.8%) and could not be amplified. Alternatively, the *E. coli* codon optimized SMT15^316-1441^ protein (∼114 kDa) containing the sulfate_transp (pfam00916), STAS domain (pfam01740), and CAP_ED domain (pfam00027) was synthesized and overexpressed in *E. coli* cells for sulfate uptake assay. Arabidopsis sulfate transporter AtSULTR1;1 (Shibagaki and Grossman, 2004) was used as a positive control. Expression of AtSULTR1;1 and SMT15^316-1441^ protein was verified by immunoblotting (Figure 1). Even though AtSULTR1;1 was predicted to be a ∼72 kDa protein, it migrated faster than predicted and this is consistent with previous observation (Shibagaki and Grossman, 2004). As expected, the cells expressing AtSULTR1;1 were capable of transporting sulfate as a substantial amount ^35^S-sulfate was detected (Table 1). The cells expressing SMT15^316-1441^ were also found to transport sulfate, supporting its sulfate transporter activity. The relatively low ^35^S-sulfate uptake for cells expressing SMT15^316-1441^ protein is likely due to a lower amount of induced SMT15^316-1441^ protein than AtSULTR1;1 protein (Figure 1).

**Figure 1.**
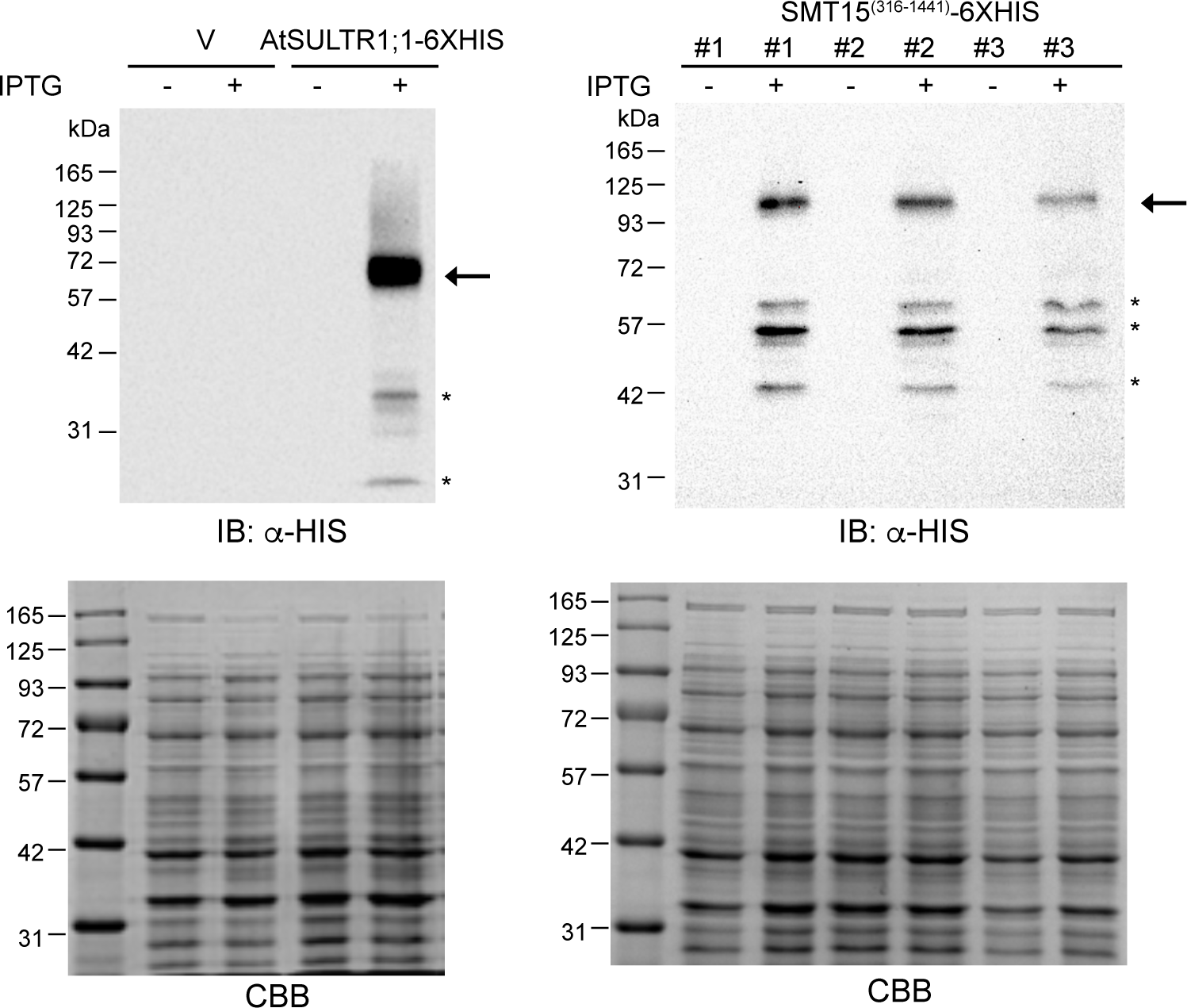
Establishment of sulfate uptake assay. Expression of *E. coli* codon-optimized SMT15^316-1441^ protein was confirmed by immunoblotting. SMT15^316-1441^ protein is shown as a single band predicted at the molecular mass ∼114 KDa. Numbers 1, 2, and 3 represent independent *E. coli* clones used for protein induction. Expression of ∼70 KDa Arabidopsis sulfate transporter (AtSULTR1:1) was also confirmed. IPTG was used to induce protein expression. Protein samples are separated by 8% SDS-PAGE. Coomassie Brilliant Blue (CBB) staining shows equal loading of total protein amount. The arrow indicates the protein at the predicted size. “*” indicates degraded protein. V, vector only control.

**Table 1.** ^35^S-sulfate uptake assay. Standard errors were derived from three independent experiments for vector control and AtSULTR1;1 strains and five independent experiments for SMT15^316-1441^ strain. T0, before incubation with ^35^S-sulfate. T10, 10 min after incubation with ^35^S-sulfate. ^a^, indicates a significant increase in sulfate uptake of AtSULT1;1 and SMT15^316-1441^ strains compared with vector control strain by a two-tailed nonparametric t test (*p* < 0.05).

| Genotype | CPM (Counts per minute) |
| --- | --- |
| <u>T0</u> |  |
| vector | 154 ± 32 |
| AtSULTR1;1 | 161 ± 35 |
| SMT15<sup>316-1441</sup> | 123 ± 11 |
| <u>T10</u> |  |
| vector | 10541 ± 745 |
| AtSULTR1;1 | 24363 ± 3455<sup>a</sup> |
| SMT15<sup>316-1441</sup> | 18724 ± 1481<sup>a</sup> |

### SMT15 protein targets to the chloroplast

To investigate where SMT15 protein is delivered to exert its function, we attempted but failed to generate a HA tagged full-length *SMT15* cDNA construct for localization study. As an alternative, constructs that allowed expression of different N-terminal fragments of SMT15 protein C-terminally fused with an enhancer YFP reporter (eYFP) were generated. Expression of the SMT15-eYFP reporters was validated by immunoblotting (Figure 2A). As expected, yellow florescence of the eYFP reporter was detected in both the cytosol and chloroplast (Figure 2B). The SMT15^N20^:eYFP reporter was also detected in both the cytosol and chloroplast (Figure 2B). As the N-terminal protein sequence of SMT15 extended to 46 amino acids, the SMT15^N46^:eYFP protein was found to target to the chloroplast (Figure 2B), indicating that the first 46 aa of SMT15 is sufficient to guide the protein into the chloroplast. Consistently, the first 63 aa of SMT15 protein also guided the eYFP reporter (SMT15^N63^:eYFP) to the chloroplast (Figure 2B). Similar results were obtained from the independent transgenic lines expressing the SMT15-eYFP reporters (Supplemental Figure S1A).

**Figure 2.**
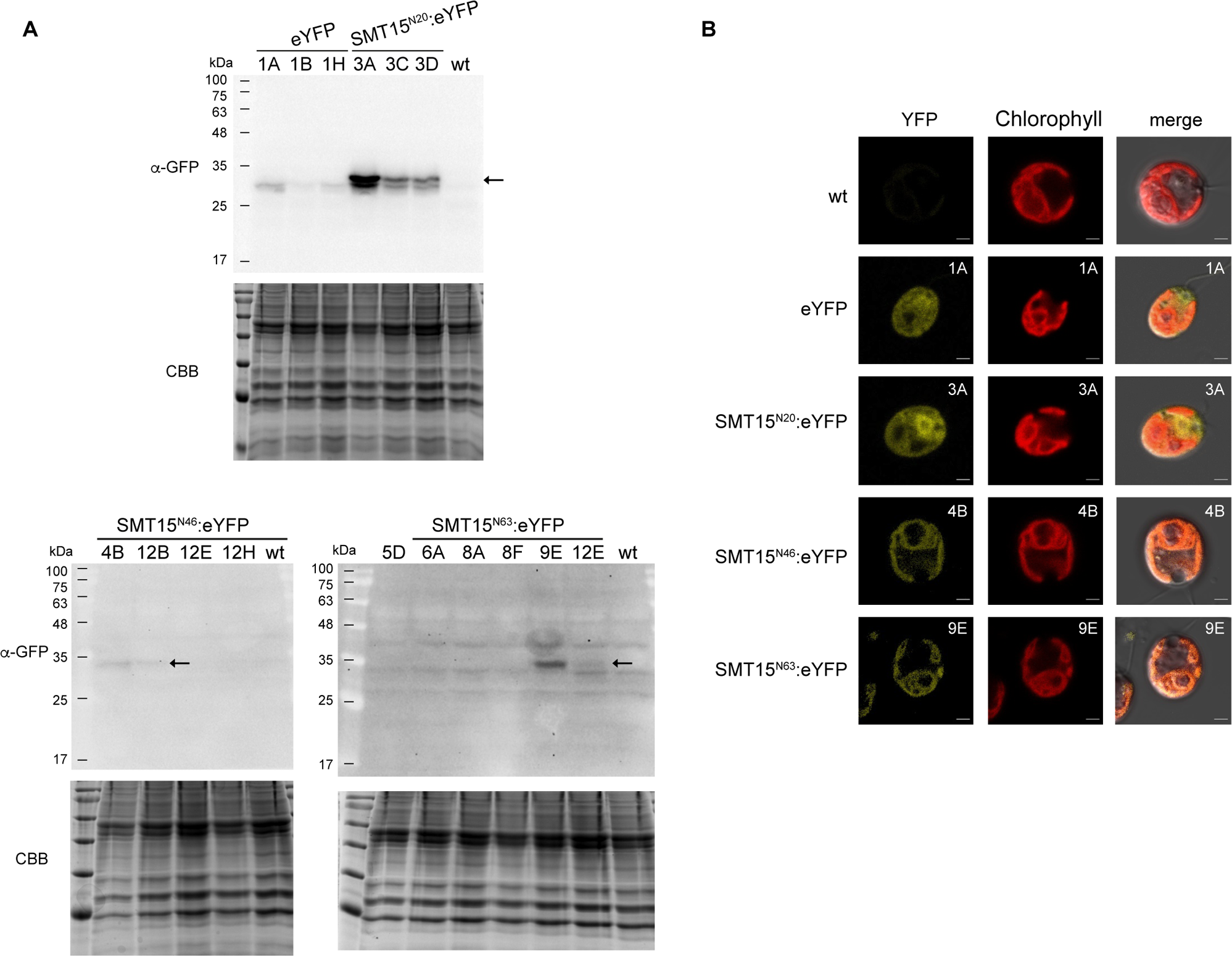
SMT15 targets the chloroplast. **A)** Expression of the indicated SMT15-eYFP reporters were confirmed by immunoblotting. Approximately 100 µg protein of each sample was separated by 12% SDS-PAGE. eYFP is ∼ 27 kDa. SMT15^N20^:YFP is ∼ 29 kDa. SMT15^N46^:YFP is ∼ 31 kDa. SMT15^N63^:YFP is ∼ 33.3 kDa. Arrow indicates the size of the indicated protein. Coomassie Brilliant Blue (CBB) staining showing equal loading. The independent transgenic lines expressing eYFP (1A, 1B, and 1H), SMT15^N20^:YFP (3A, 3C, and 3D), SMT15^N46^:YFP (4B, 12B, 12E, and 12H), SMT15^N63^:YFP (5D, 6A, 8A, 8F, 9E, and 12E) are shown. **B)** Confocal images showing subcellular localization of the indicated SMT15-eYFP reporters. wt, wild-type. Chlorophyll, chlorophyll autofluorescence in the RFP channel. DIC, differential interference contrast images of cells superimposed with YFP and chlorophyll signals. Scale bar, 2 μM.

To examine subcellular localization of SMT15, a SMT15-specific antibody was generated. However, this SMT15 specific antibody recognized multiple non-specific bands in the total cellular extract in immunoblotting and was therefore omitted from immunolocalization analysis. Even so, the SMT15-specific antibody was able to detect a ∼144 kDa SMT15 protein in membrane factions prepared from the wild-type and complemented *smt15-1 pSMT15.1 #7* strains but not from the *smt15-1* strain (Supplemental Figure S1B), confirming that SMT15 is a membrane protein (Fang et al., 2014). Taking these results together, we concluded that SMT15 is a chloroplast membrane-associated sulfate transporter.

### Decreasing glutathione levels in *smt15-1* strain restores its cell-size defect

Because elevated glutathione content is co-segregated with the small cell-size of *smt15-1* (Fang et al., 2014), we were interested in testing whether the increased glutathione caused increased cell division number and therefore generation of small sized daughters in the *smt15-1* mutant. To test this idea, artificial microRNA (amiRNA) constructs targeting two different loci of 3′ UTR of *γ-GLUTAMYLCYSTEINE SYNTHETASE* (*GSH1*, the rate-limiting enzyme required for glutathione biosynthesis) were generated (amiGSH1-1 and amiGSH1-2) and introduced into the *smt15-1* mutant (Figure 3A). Even though multiple transgenic strains *smt15-1::amiGSH1-*1 were obtained, expression of *GSH1* was not affected or mildly reduced (Figuyre 3B). The *smt15-1::amiGSH1-*1 transgenic lines were omitted from further analysis. Downregulation of *GSH1*, on the other hand, was validated in five independent *smt15-1::amiGSH1-*2 strains (Figure 3B). We chose to investigate the *smt15-1::amiGSH1-2* #3B, #4D, and #4F lines because their *GSH1* mRNA was approximately 30% or less than that of *smt15-1* mutant (Figure 3B). Decrease in total glutathione level was confirmed in the *smt15-1::amiGSH1-2* #4D and #4F lines (Figure 3C). Importantly, decreased glutathione level increased the cell-size of the *smt15-1* cells to a size similar to wild-type cells (Table 2). Even though glutathione level did not show significant reduction in *smt15-1::amiGSH1-2* #3B strain, it generated daughters larger than that of *smt15-1* strain (Table 2). Hence, we conclude that defective glutathione homeostasis in *smt15-1* mutant caused increased cell division number and small cell-size.

**Figure 3.**
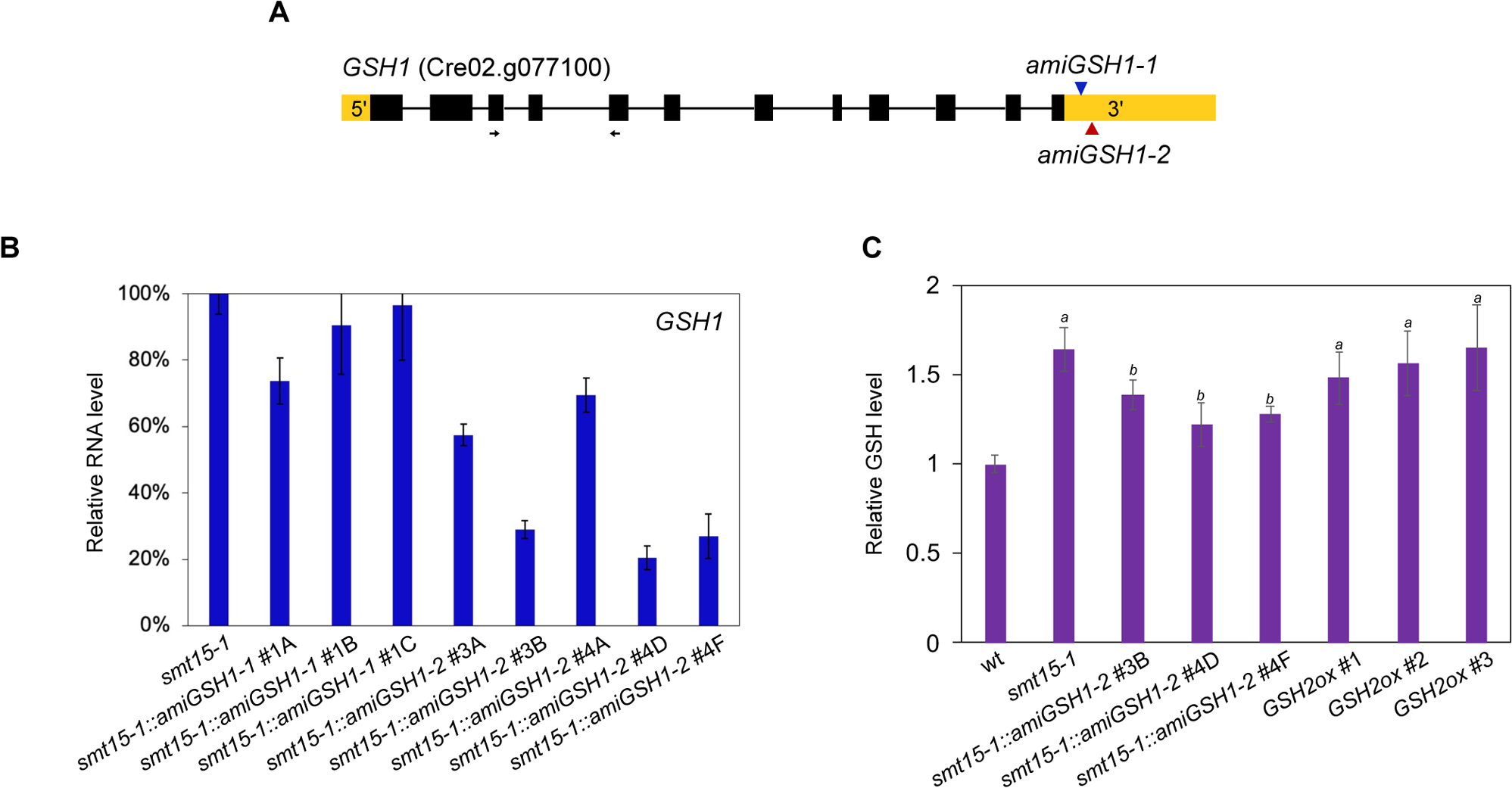
Generation of the *GSH1* artificial microRNA lines. **A)** Schematic of the GSH1 genome DNA structure. Black boxes indicate exons. Black lines indicate introns. Yellow boxes indicate 5’ or 3’ UTR regions. The blue triangle indicates the 3’ UTR position where amiGSH1-1 was targeted. The red triangle indicates the 3’ UTR position where amiGSH1-2 was targeted. The arrows indicate primers used for RT-qPCR. **B)** Relative expression level of *GSH1* in three independent *smt15-1::amiGSH1-1* lines and five independent *smt15-1::amiGSH1-2* were analyzed by RT-qPCR. **C)** Relative GSH content of three independent *smt15-1::amiGSH1-*2 lines and three independent *GSH2ox* lines. Standard errors were derived from five biological replicates. The glutathione level of wt cells was arbitrarily set to one. *a*, indicates a significant increase in cell size compared with wt cells by a two-tailed t test (*p* < 0.05). *b*, indicates a significant decrease in cell size compared with *smt15-1* cells by a two-tailed t test (*p* < 0.05).

**Table 2.** Daughter cell size distribution of the *smt15-1::amiGSH1* strains. Standard errors were derived from three biological replicates. *a*, indicates a significant increase in cell size compared with *smt15-1* cells by a two-tailed t test (*p* < 0.05). *b*, indicates a significant decrease in cell size compared with wt cells by a two-tailed t test (*p* < 0.05).

| Genotype | Daughter cell size<br>($\mu\text{m}^3$) |
| --- | --- |
| wild-type | $64.0 \pm 0.8$ |
| *smt15-1* | $52.6 \pm 1.3^b$ |
| *smt15-1::amiGSH1-2* #3B | $62.1 \pm 0.4^a$ |
| *smt15-1::amiGSH1-2* #4D | $62.9 \pm 1.0^a$ |
| *smt15-1::amiGSH1-2* #4F | $60.9 \pm 1.5^a$ |

### Glutathione is important to regulate size-mediated cell division

To directly test whether glutathione regulates cell division in Chlamydomonas, cell cycle progression was monitored in cells treated with diethyl maleate (DEM), which depletes glutathione by direct conjugation (Plummer et al., 1981). DEM has been commonly used to study glutathione and cell cycle regulation in animal and plant systems (Markovic et al., 2009; Diaz Vivancos et al., 2010; Jiang et al., 2015). DEM at the concentration of 0.5 mM, 1 mM, or 2 mM was added into the dark-incubated unsynchronized cultures (G_0_) one hour before switching to light (at the time point of T-1, Figure 4A) to minimize cell cycle-associated glutathione fluctuation (Fang et al., 2014). DEM treatment at a concentration of 1 mM or 2 mM caused a substantial decrease in glutathione level at 5 to 9 h (4 to 8 h after switching to light, T4 to T8) after treatment (Figure 4B). DEM at the concentration of 0.5 mM caused a slight increase in total glutathione. It is possible that glutathione depletion by 0.5 mM DEM triggers *de novo* glutathione biosynthesis as a compensatory mechanism to replenish glutathione. Even so, 0.5 mM DEM treatment delayed cell entry to mitosis (Figure 4C). Moreover, glutathione depletion by 1 mM or 2 mM DEM completely blocked cell cycle progression (Figure 4C) and slowed down cell growth (Figure 4D), indicating the importance of glutathione for cell growth and division. To investigate whether increased glutathione level affects cell division, glutathione or glutathione ethyl ester was added to wild-type cells. However, adding exogenous glutathione or glutathione ethyl ester did not increase cellular glutathione and was therefore not pursued further.

**Figure 4.**
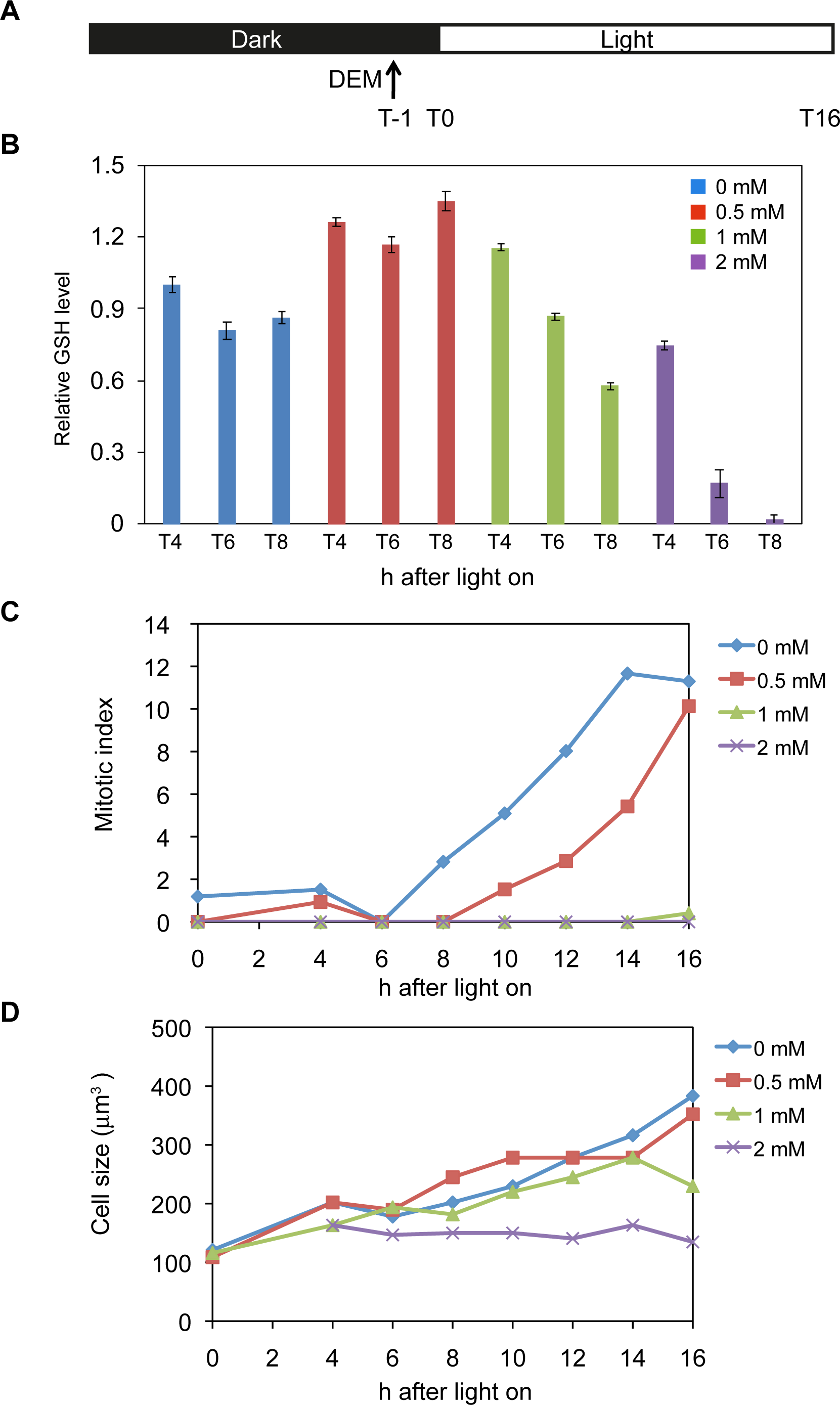
Diethyl maleate (DEM) treatment affects cell growth and entry into the cell cycle in the unsynchronized Chlamydomonas cultures. **A)** Graph depicting the design of the experiment showing when DEM was added. **B)** Glutathione levels of cultures sampled at the indicated number of hours after switching to light upon DEM treatment. The total glutathione content was measured using a Glutathione Detection Kit (Enzo Life Sciences) following the manufacturer’s instructions as described previously (Fang et al., 2014). **C)** Mitotic index of DEM-treated cells after switching to light. **D)** Modal cell size of DEM-treated cells after switching to light.

To increase cellular glutathione in wild-type cells, alternatively we generated the transgenic lines overexpressing the *γ-GLUTAMYLCYSTEIN LIGASE* (*GSH1*) or *GLUTATHIONE SYNTHETASE* (*GSH2*) that encodes the first or second catalytic enzyme required for glutathione biosynthesis. Overexpression of *GSH1* and *GSH2* has been reported to increase cellular glutathione level in plants (Noctor et al., 1996; Noctor et al., 1998; Creissen et al., 1999; Zhu et al., 1999). Multiple transgenic lines carrying the pF1-GSH1-3XFLAG construct were generated. Unfortunately, none of the *GSH1ox* lines expressed the GSH1 protein and were omitted from further analysis. On the other hand, expression of GSH2 protein was confirmed in *GSH2ox* #1*, GSH2ox* #2, and *GSH2ox* #3 strains carrying the GSH2-3XFLAG construct (Figure 5A). Overexpression of GSH2 was shown to increase glutathione in *GSH2ox* #1*, GSH2ox* #2, and *GSH2ox* #3 strains (Figure 3C). Importantly, all of the *GSH2ox* transgenic lines generated small sized daughters that resembled those of *smt15-1* mutant (Figure 5B and Table 3), indicating that glutathione is sufficient to increase cell division number.

**Figure 5.**
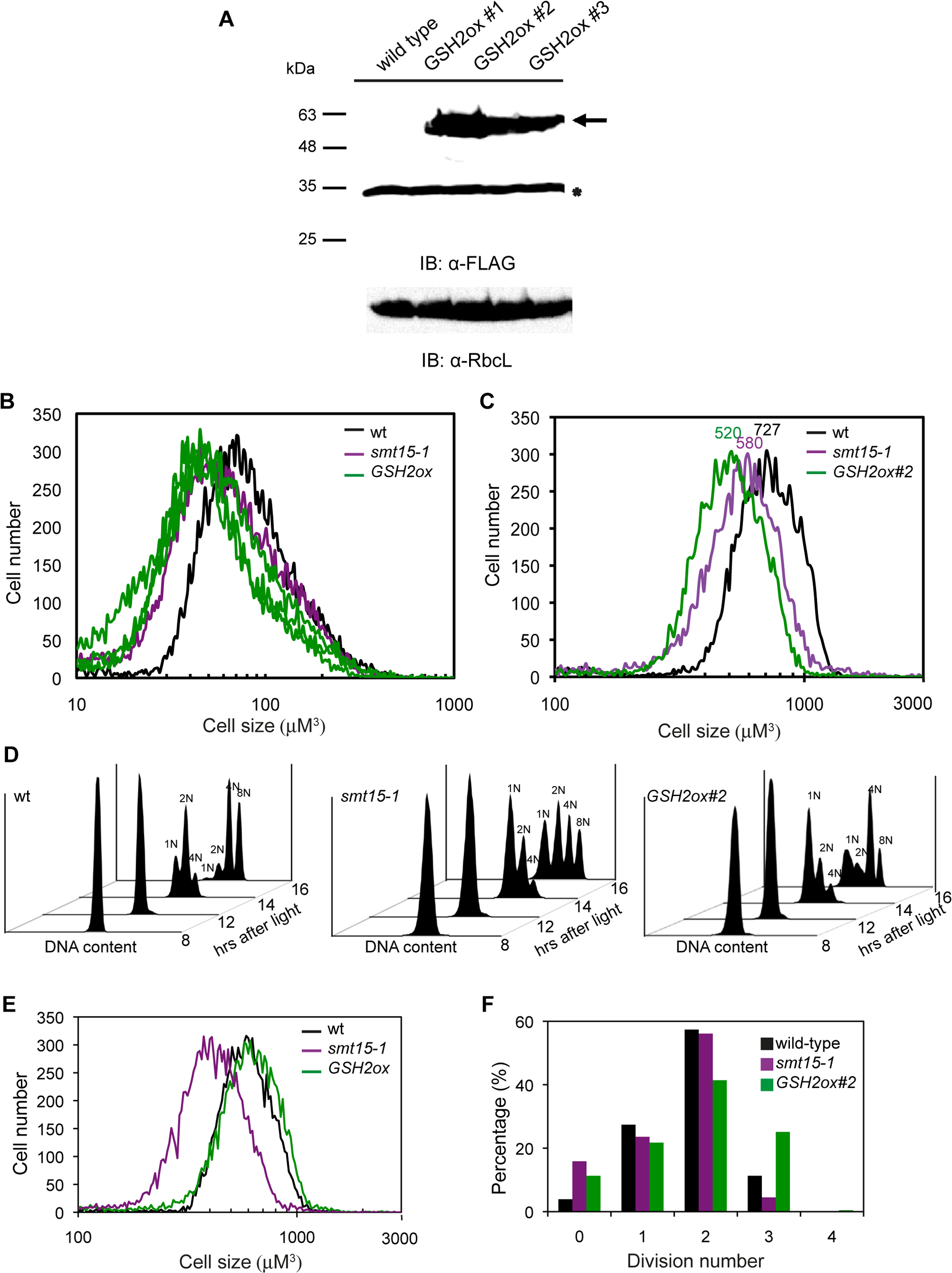
Overexpression of GSH2 results in increased cell division number. **A)** Immunoblotting showing expression of GSH2 protein in three independent transgenic lines. As a loading control, the blot was stripped and reprobed with RbcL antibodies. **B)** Size distributions of dark-shifted unsynchronized cells from *smt15-1* and three independent *GSH2* overexpressors (*GSH2ox*, in green lines). **C)** Size distributions of light-grown cells sampled at 14 h after light from the synchronized wild-type, *smt15-1* and *GSH2ox* #2 cultures. Number indicated on the top of the size distribution plot represents modal cell size of the indicated strain. **D)** FACS profile showing the DNA content of the synchronized wild-type, *smt15-1*, and *GSH2ox* #2 cultures during cell cycle progression. DNA content was indicated above each peak at 14 and 16 h after light. The number of divisions was confirmed by microscopic examination. **E)** Size distribution of the synchronized wild-type and *smt15-1*, and *GSH2ox* #2 cultures collected at 14 h after light and before dark shift. **F)** Distribution of cell division numbers from 14 h light-grown synchronized cells of the indicated genotype.

**Table 3.** Daughter cell size distribution, of the *GSH2* overexpressing strains and indicated strains. Standard errors were derived from at least three biological replicates. *a*, indicates a significant decrease in cell size compared with wt cells by a two-tailed t test (*p* < 0.05).

| Genotype | Daughter cell size<br>($\mu\text{m}^3$) |
| --- | --- |
| wild-type | $68.3 \pm 2.2$ |
| *smt15-1* | $48.7 \pm 3.1^a$ |
| *GSH2ox* #1 | $47.5 \pm 2.1^a$ |
| *GSH2ox* #2 | $49.3 \pm 2.5^a$ |
| *GSH2ox* #3 | $50.2 \pm 2.9^a$ |

To validate the increase in cell division number and generation of small sized daughters in *GSH2ox* strains by glutathione, wild-type, *smt15-1*, and *GSH2ox* #2 transgenic strains were synchronized using 14-h-light/10-h-dark cycles and their cell division was examined by DNA content as cells pass S/M phase using image cytometry. For wild-type cells, the modal cell size, the size occurred most often in the dataset (Fang et al., 2006), of cells growing photosynthetically under light for 14 h reached up to ∼700 μm^3^ (Figure 5C) and the mother cells were able to divide up to three times in the dark (8N, Figure 5D). Importantly even though the modal cell size of *smt15-1* and *GSH2ox* #2 cultures only reached up to ∼580 μm^3^ and 520 μm^3^ respectively in the light (Figure 5C), they were also capable of dividing up to three times in the dark (8N, Figure 5D). Moreover, a very small fraction of the *smt15-1* mother cells that were capable of dividing four times was observed.

To validate the image cytometry data, division number was measured by examining division clusters of cells grown for 14 h as described previously (Fang et al., 2006). Even though the mother cells of wild-type and *GSH2ox* #2 cells shared similar size distribution this time (Figure 5E), more than 25% of *GSH2ox* #2 cells divided three times (8-cell clusters) while only 11% of wild-type cells could form 8-cell clusters (Figure 5F). Moreover, a small fraction of *GSH2ox#2* cells (∼0.4%) was capable of dividing four times and formed 16-cell clusters. Because *GSH2ox* #3 could not be synchronized by 14-h-light/10-h-dark cycles, 12-h-light/12-h-dark cycles were used instead. Consistent with the results from *GSH2ox* #2 and *smt15-1* strains, the synchronized *GSH2ox* #3 cells were able to divide up to three times or even four times despite their relatively small-sized mothers (Supplemental Figure S2A) while the vast majority of the wild-type mothers only divided twice (Supplemental Figure S2B). Together, our results demonstrated that similar to *smt15-1,* increased cell division number of the *GSH2ox* transgenic lines enabled generation of small-sized daughters.

### Enhanced basal body-associated glutathione at early S/M phase is associated with increased cell division in *smt15-1* and *GSH2ox* lines

Because both *smt15-1* and *GSH2ox* strains had increased cell division number and generated small daughter cells, we speculated that cell cycle-associated subcellular accumulation of glutathione may be important to regulate size-mediated cell division. To test this idea, we used 5-chloromethylfluorescein diacetate (CMFDA) to monitor subcellular glutathione in the synchronized wt and *smt15-1*. CMFDA has been frequently used to monitor glutathione in both animal and plant systems (Voehringer et al., 1998; Lantz et al., 2001; Markovic et al., 2007; Diaz Vivancos et al., 2010). Synchrony of wt and *smt15-1* cultures were confirmed by mitotic index analysis (Figure 6A) and expression of the S/M-specific markers (Figure 6B), *CYCLIN-DEPENDENT KINASE B1* (*CDKB1*) and *PROLIFERATING CELL NUCLEAR ANTIGEN* (*PCNA*). During the G1 phase (6 h after light incubation), glutathione appeared to be evenly distributed in the cytosol of wt and *smt15-1* cells (Figure 6C). The glutathione signal reduced substantially in wt cells as cells entered the S/M phase (12 h after light incubation). This result is consistent with our previous study in which we reported that glutathione level is reduced during the S/M phase (Fang et al., 2014). Even though glutathione was reduced in the cytosol of *smt15-1* cells as cells entered the S/M phase, glutathione was found to be specifically concentrated in the basal bodies in more than 70% of *smt15-1* cells (Figure 6C and Table 4). This experiment was repeated two more times and similar results were obtained (Supplemental Figure S3). Hence, our results revealed that glutathione was enriched in the basal bodies of *smt15-1* cells at the S/M phase.

**Figure 6.**
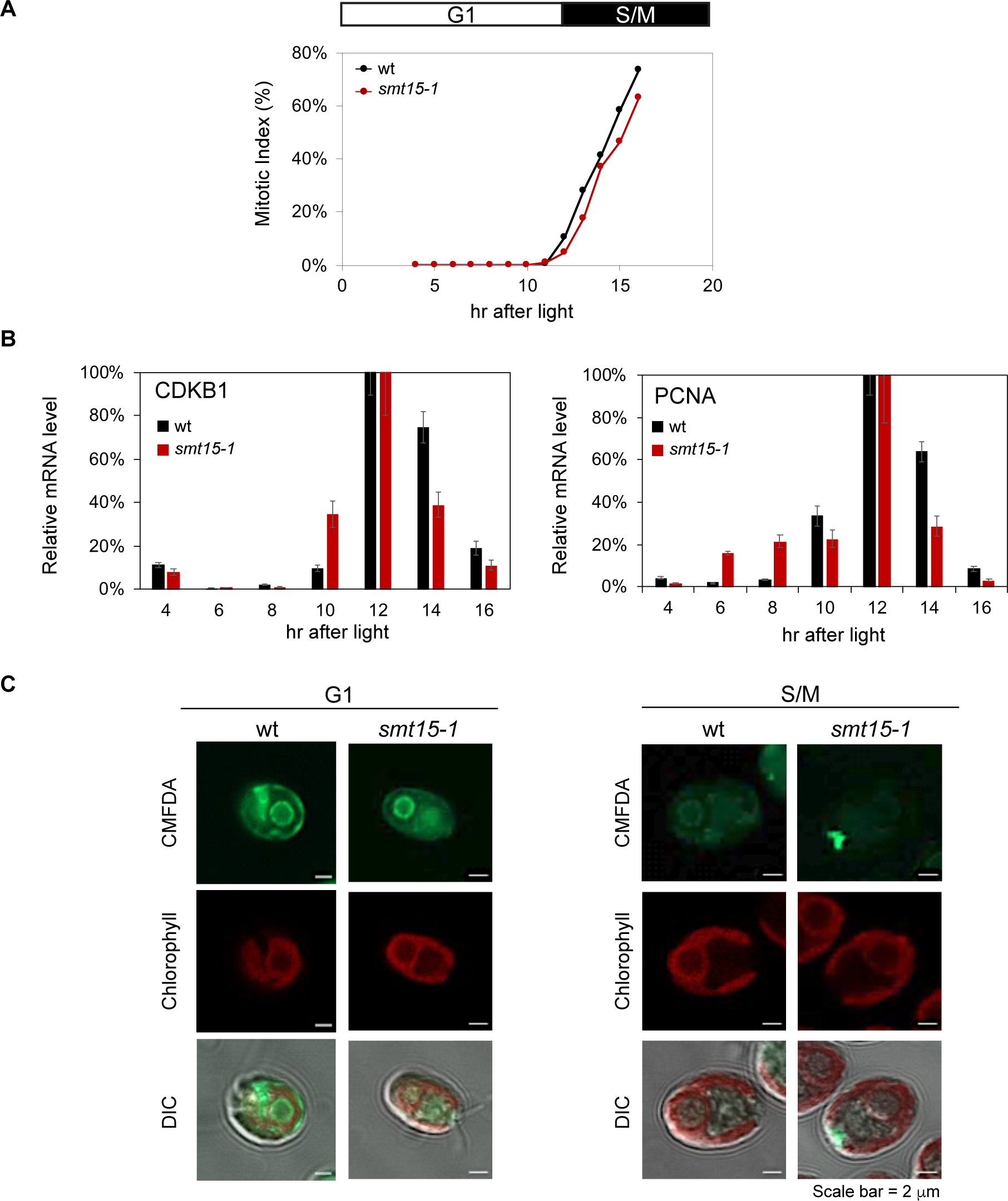
Subcellular localization of glutathione in wild-type and *smt15-1* cells at G1 and S/M phases. **A)** Graph showing mitotic index of the synchronous cultures. Wild-type (wt) and *smt15-1* cultures entered S/M phase at approximately 12 h. The cell cycle phases are indicated by bars above the graph. The wt strain is labeled in black and *smt15-1* is labeled in red. Synchronized cultures were maintained in 12-h-light/12-h-dark cycles. **B)** Expression of S/M phase markers *CDKB1* and *PCNA* determined by RT-qPCR, with SE shown by error bars. C, CMFDA staining of G1 (collected at 6 h after light) and S/M phase (collected at 12 h after light) of wt and *smt15-1* cells. Chlorophyll, chlorophyll autofluorescence. DIC, differential interference contrast images of cells superimposed with CMFDA and chlorophyll signals.

**Table 4.**
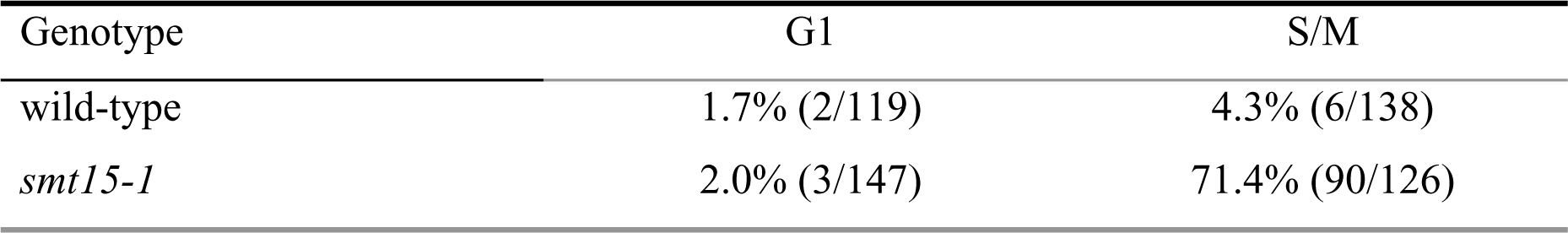
Basal body-associated glutathione at G1 and early S/M phases examined by CMFDA staining. Percentage was calculated by number of cells shown basal-body-associated glutathione was divided by the number of examined cells.

To test whether glutathione concentrated in the basal bodies is relevant to the increased cell division number that generates small sized daughters in *GSH2* overexpressors, we monitored glutathione of the synchronized *GSH2ox* #2 and *GSH2ox* #3 cell cultures. Synchrony of wt, *GSH2ox* #2, and *GSH2ox* #3 strains were confirmed by mitotic index analysis (Figure 7A and Supplemental Figure S4A) and expression of the S/M-specific markers *CDKB1* and *PCNA* (Figure 7B and Supplemental Figure S4B). Cellular glutathione of wt, *GSH2ox* #2, and *GSH2ox* #3 strains at G1 phase (6 h after light incubation) and S/M phase (12 h after light incubation) was monitored by CMFDA staining. Similar to wt cells, glutathione appeared to localize primarily in the cytosol in *GSH2ox* #2, and *GSH2ox* #3 strains at G1 phase (Figure 7C and Supplemental Figure S4C). However, unlike wt cells whose cytosolic glutathione was homogenous, we found glutathione was sometimes concentrated in a few foci of the cytosol in *GSH2ox* lines (Figure 7C). Regardless of whether the cells were *GSH2ox* #2 or *GSH2ox* #3 strain, cytosolic glutathione was reduced substantially as cells entered S/M phase similar to wt and *smt15-1* cells. Importantly, the basal body-specific enrichment of glutathione was also detected in more than 80% of *GSH2ox* #2 and *GSH2ox* #3 cells entering S/M phase (Table 5), suggesting increased glutathione in the basal body is linked to the increased cell division number in *smt15-1* and *GSH2ox* transgenic lines.

**Figure 7.**
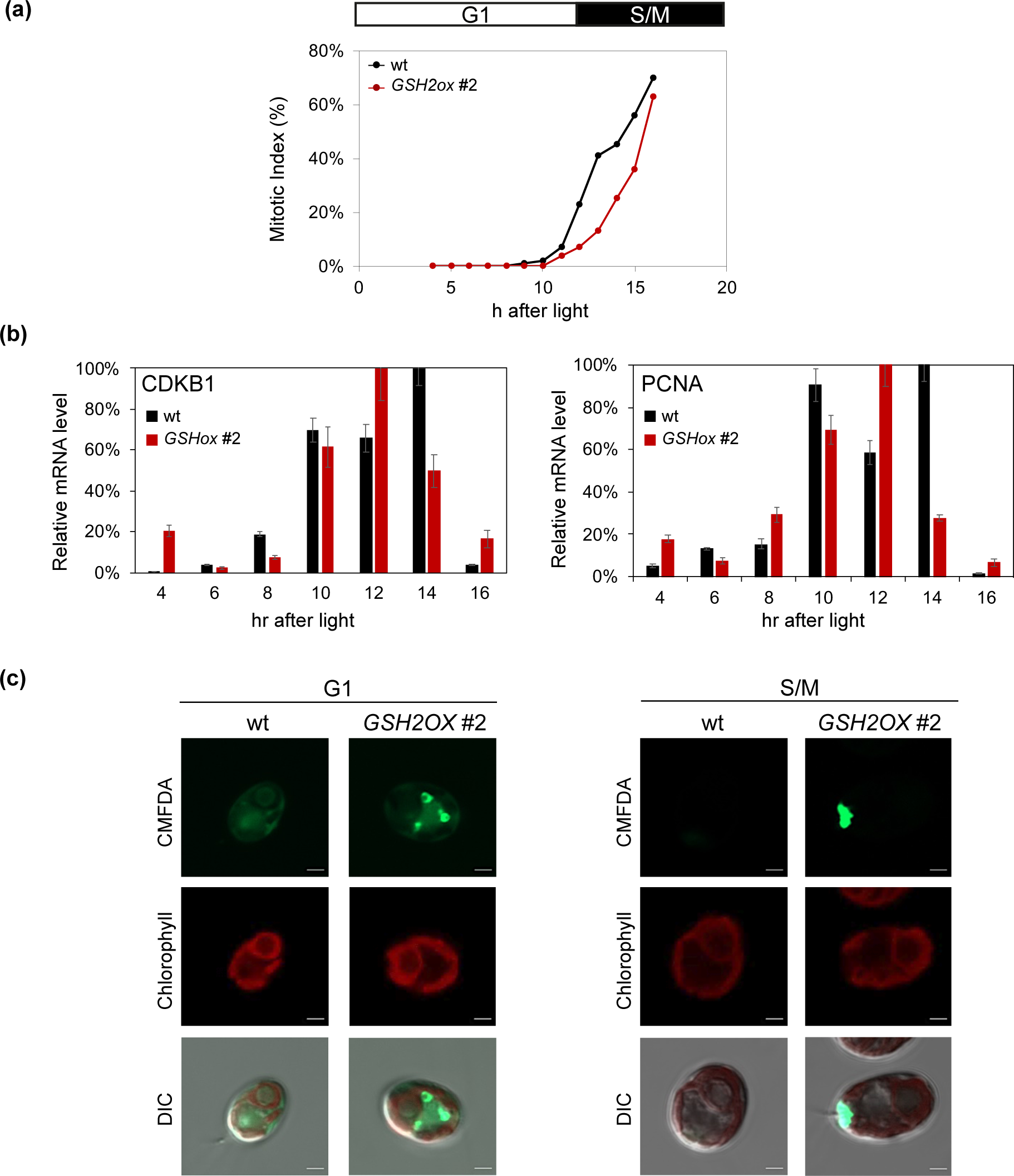
Subcellular localization of glutathione in wild-type and *GSH2ox* #2 cells at G1 and S/M phases. **A)** Graph showing mitotic index of the synchronous cultures. Wild-type (wt) and *GSH2ox* #2 cultures entered S/M phase at approximately 12 h. The cell cycle phases are indicated by bars above the graph. The wt strain is labeled in black and *GSH2ox* #2 is labeled in red. Synchronized cultures were maintained in 12-h-light/12-h-dark cycles. **B)** Expression of S/M phase markers *CDKB1* and *PCNA* determined by RT-qPCR, with SE shown by error bars. C, CMFDA staining of G1 (collected at 6 h after light) and S/M phase (collected at 12 h after light) of wt and *GSH2ox* #2 cells. Chlorophyll, chlorophyll autofluorescence. DIC, differential interference contrast images of cells superimposed with CMFDA and chlorophyll signals. Scale bar, 2 μM.

**Table 5.** Basal body-associated glutathione at G1 and early S/M phase examined by CMFDA staining. The percentage was calculated by the number of cells showing basal-body-associated glutathione divided by the number of examined cells. ^a^, CMFDA staining of the tested cells is shown in Figure 7. ^b^, CMFDA staining of the tested cells is shown in Figure S4.

| Genotype | G1 | S/M |
| --- | --- | --- |
| wild-type<sup>a</sup> | 2.7 % (3/113) | 6.8 % (10/146) |
| *GSH2ox* #2<sup>a</sup> | 1.4 % (2/138) | 81.8 % (117/143) |
| wild-type<sup>b</sup> | 2.0 % (2/111) | 0 % (0/140) |
| *GSH2ox* #3<sup>b</sup> | 0 % (0/125) | 63.2% (91/144) |

### Glutathione is mainly synthesized in the cytosol

To investigate how a chloroplast-associated sulfate transporter SMT15 regulates cellular glutathione homeostasis, it is necessary to know the subcellular compartment(s) in which glutathione is synthesized. In mammals and budding yeast, glutathione is synthesized in the cytosol by two cytosolic enzymes, λ-glutamylcysteine synthetase (GSH1) and glutathione synthetase (GSH2) (Griffith and Meister, 1985; Toledano et al., 2013). In Arabidopsis, the cytosolic and plastidic dual targeting GSH2 protein enables glutathione to be synthesized in the cytosol and the chloroplasts (Wachter et al., 2005; Pasternak et al., 2008). We reasoned that as *GSH1* and *GSH2* are single-copy genes in Chlamydomonas (Merchant et al., 2007), subcellular localization of GSH1 and GSH2 protein would provide information on where glutathione is synthesized. Even though the anti-FLAG antibody was able to successfully detect the GSH2-3XFLAG protein in *GSH2ox* transgenic lines by immunoblotting (Figure 5A), it failed to detect reliable immunofluorescence (IF) signals. To overcome this problem, a Gly-rich linker was inserted between sequences encoding GSH1 or GSH2 and 3XFLAG tag to generate the pF1-GSH1-lin-3XFLAG and pF1-GSH2-lin-3XFLAG constructs. Insertion of a flexible Gly-rich linker between a tagged protein and an epitope has been shown to increase epitope accessibility (Sabourin et al., 2007; Reddy Chichili et al., 2013) and improve IF analysis (Lin et al., 2020). Therefore, pF1-GSH1-lin-3XFLAG and pF1-GSH2-lin-3XFLAG were introduced into the *uvm4* cells, which enhances expression of transgenes (Neupert et al., 2009). Similar to our previous attempt, GSH1 protein failed to be expressed in the *GSH1-lin-3XFLAG* transgenic lines, suggesting overexpressing GSH1 protein may be cytotoxic. Alternatively, we generated N-terminal GSH1 translation reporters to investigate where GSH1 protein is targeted. To this end, cDNA sequences encoding the first 21, 30, 42, and 60 amino acids of GSH1 were translationally fused to a CrVenus reporter (Mackinder et al., 2016) and expression of the the GSH1 reporters (GSH1^n21^: Venus, GSH1^n30^:Venus, and GSH1^n42^:Venus) were verified by immunoblotting (Supplemental Figure S5A). Because GSH1^n60^:Venus protein could not be detected in the transgenic lines, it was omitted from further analysis. Regardless the lengths of N-terminal GSH1 protein introduced, CrVenus florescence was mainly detected in the cytosol (Supplemental Figure S5B), suggesting λ-glutamylcysteine is synthesized in the cytosol.

As for *GSH2*, multiple *GSH2-lin-3XFLAG* transgenic lines overexpressing the GSH2-lin-3XFLAG protein were confirmed by immunoblotting (Figure 8A). Two of the transgenic lines, *GSH2-lin-3XFLAG* #7C and *GSH2-lin-3XFLAG* #7D, were chosen for immunolocalization study because they had relatively high GSH2-3xFLAG protein level. Based on the immunofluorescence result, the GSH2-3X FLAG protein was localized predominantly in the cytosol (Figure 8B), inferring that glutathione biosynthesis occurs primarily in the cytosol in Chlamydomonas. This result coincided with the CMFDA analysis showing that glutathione is accumulated primarily in the cytosol (Figure 6C and Figure 7C).

**Figure 8.**
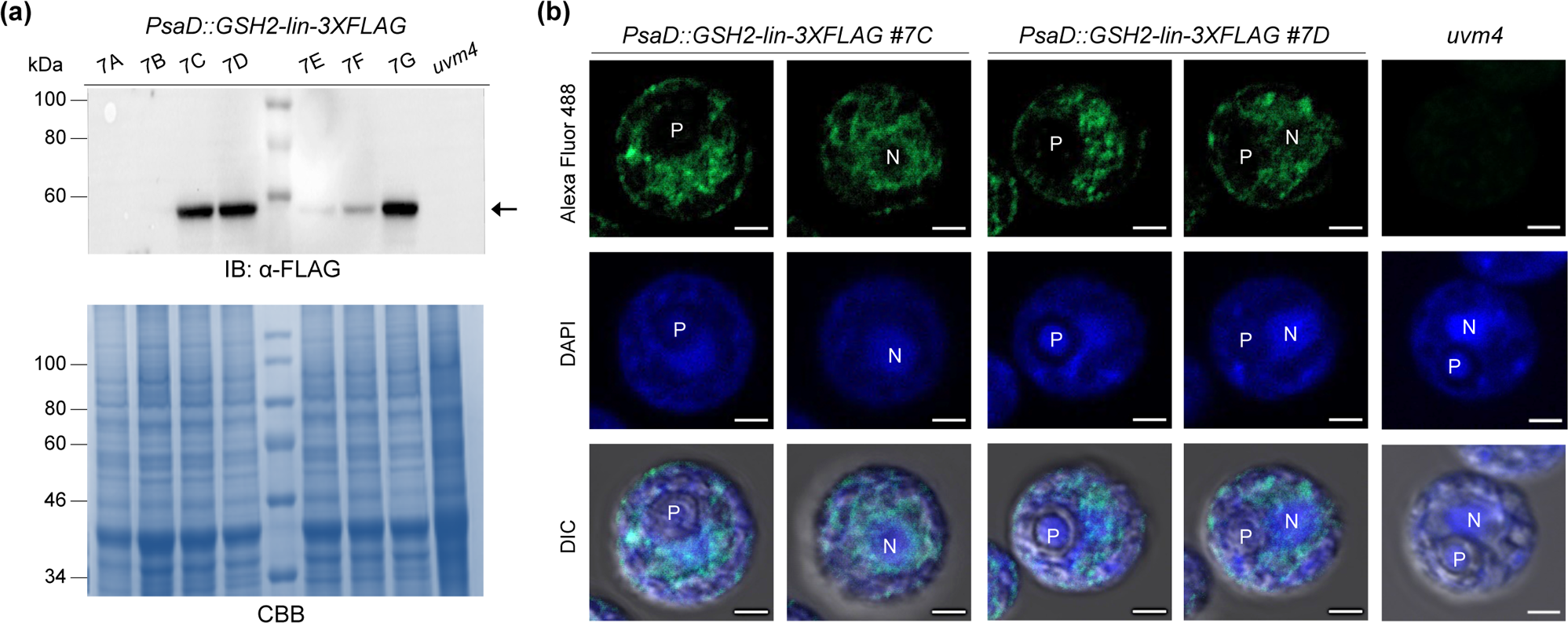
Subcellular localization of the GSH2 protein. **A)** Immunoblotting showing expression of GSH2 protein in seven independent transgenic lines in the *uvm4* background. The arrow indicates expression of GSH2-lin-3XFLAG at the predicted size of 56.5 kDa. Coomassie brilliant blue (CBB) staining is used to show equal loading of protein per lane. **B)** Immunolocalization of the GSH2 protein in *PsaD::GSH2-lin-3XFLAG* #7C and *PsaD::GSH2-lin-3XFLAG* #7D transgenic lines. *uvm4* strain was used as a negative control. P, pyrenoid. N, nucleus. Scale bar, 2 μM.

## Discussion

Glutathione is important for cell cycle regulation in plant and animal systems (Pallardó et al., 2009; Diaz Vivancos et al., 2010; García-Giménez et al., 2013; Diaz-Vivancos et al., 2015). We previously isolated a small cell-size mutant, *smt15-1*, whose sulfur acclimation response and cell-cycle associated glutathione dynamics are altered (Fang and Umen, 2008; Fang et al., 2014). Here, we demonstrated that *SMT15* encodes a *bona fide* chloroplast sulfate transporter and its function is important in the maintenance of cellular glutathione homeostasis. Importantly, decreasing glutathione levels in the *smt15-1* mutant generated daughters with a cell-size similar to those of the wt strain. Consistent with this notion, overexpression of *GSH2* increased cell division number and generated small cells that recapitulated the cell size of *smt15-1* mutant. Together, our data provided the evidence linking glutathione to the cell division cycle in Chlamydomonas.

During the course of our study, we discovered that glutathione, which is mainly synthesized in the cytosol (Figure 8), was accumulated in relatively high abundance in the cytosol during the G1 phase and gradually decreased as cells went into S/M phase in wt cells (Figure 6C). Similar to wt cells, glutathione was accumulated in the cytosol of *smt15-1* cells during G1 phase and decreased in abundance as cells entered S/M phase (Figure 6C). Surprisingly for *smt15-1* cells at the S/M phase, glutathione was concentrated in the basal bodies (Figure 6C). This result helps explain why total glutathione of *smt15-1* cells fails to decline during the S/M phase (Fang et al., 2014). Together, our data showed that glutathione is temporally and spatially regulated during the cell cycle in Chlamydomonas. The question then becomes, why decreasing the cytosolic glutathione would coincide with entry into S/M phase. Even though it is not clear that downregulation of cytosolic glutathione is relevant to cell cycle control in Chlamydomonas, the redox cycle is reported to coordinate with the cell cycle in mammalian cells and budding yeast (Tu et al., 2005; Hoffman et al., 2008). Moreover, the oxidative cellular environment appears to initiate cell division in mammalian cells (Menon and Goswami, 2007; Burhans and Heintz, 2009). Coincidently, thiol oxidation of overall proteins is reported to increase during mitosis (Patterson et al., 2019). Hence, we speculated that a decrease in cytosolic glutathione may establish a relatively oxidized environment, which prepares cells to enter S/M phase in Chlamydomonas. Whether thiol-dependent redox regulates cell cycle progression remains to be determined.

The abundant basal body-associated glutathione detected at early S/M phase in both *smt15-1* and GSH2 overexpressing strains was quite unexpected. It is not clear how glutathione is transported and concentrated in the basal bodies at S/M phase. Nevertheless, the positive link between elevated glutathione in the basal body and increased number of cell divisions suggests that increased glutathione level in the basal body at S/M phase may facilitate cell division that leads to generation of small daughters in the *smt15-1* strain and GSH2 overexpressing lines. It is unclear exactly how the redox status of the basal body affects cell division. In Eukaryotes, basal bodies and their structural equivalents, centrioles, organize microtubules in the cytoplasm to form the spindle apparatus that is important for cell division and assembly of flagella or cilia (Dutcher, 2003; O’Toole and Dutcher, 2014). Basal body replication is coupled with cell division and occurs during the prophase. Because basal body is required for coordination of mitosis and cytokinesis, defects in genes involved in basal body formation often have a defect in cell division (Ehler et al., 1995; Dutcher and Trabuco, 1998; Silflow et al., 2001; Matsuura et al., 2004; Keller et al., 2010; Goyal et al., 2014). Even though it is unclear whether glutathione regulates the redox environment within the basal body that subsequently modulates its functions, recent redox regulation studies in the basal body/centrosome have revealed some insights. For example, reversible oxidative inhibition of the centrosome-bound Cdc14B tyrosine phosphatase, which is essential for mitosis exit, facilitates CDK1-dependent mitosis process (Lim et al., 2015). In addition, oxidation of a conserved cysteine residue within Aurora A kinase (AURKA) protein that is required for centrosome maturation and progression of mitosis (Willems et al., 2018) promotes its activation during mitosis (Lim et al., 2020; Lim et al., 2020). Intriguingly, both Chlamydomonas AURKA and CDC14 proteins are flagella-associated proteins (Pan and Snell, 2000; Wang et al., 2014). The potential functions of Chlamydomonas AURKA and CDC14 in mitosis remain to be determined.

In addition to the basal body, cilium/flagella biogenesis also affects the cell cycle progression (Jackson, 2011; Kim et al., 2011; Li et al., 2011). Interestingly, the basal body-specific nitric oxide synthases that are capable of producing nitric oxide and superoxide regulate ciliogenesis (Xue et al., 1996; Simet et al., 2013). Additionally, incubation with N-acetyl-L-cysteine, a precursor of glutathione, prevented loss of ciliogenesis caused by virus infection in human bronchial epithelial cells (Mata et al., 2012). Interestingly, thioredoxin domain-containing proteins and peroxiredoxin-mediated redox regulation are involved in ciliogenesis (Blackburn et al., 2017; Ji et al., 2019; Price and Sisson, 2019). Hence, it is possible that thiol-mediated redox regulation in basal bodies and/or cilliogenesis has impacts on cell division cycle.

How does the chloroplast sulfate transporter SMT15 affect glutathione homeostasis in Chlamydomonas? In Chlamydomonas, sulfate is activated and reduced by the chloroplast-localized adenosine 5′-phosphosulfate reductase (APR) and sulfite reductase (SIR) to form sulfite (Patron et al., 2008). Sulfite is then combined with *O-*acetyl-Ser to form cysteine, the precursor of glutathione, in the chloroplast (Ravina et al., 2002). Considering *smt15-1* cells have elevated glutathione, it is unlikely that SMT15 acts as a chloroplast sulfate importer because reducing sulfate supply in the chloroplast would decrease cysteine synthesis that would in turn lead to reduced glutathione. Moreover, transport of the cytosolic sulfate to the chloroplast has been reported to be carried out by two chloroplast-localized sulfate permeases, SulP1 and SulP2 (Chen et al., 2003; Chen and Melis, 2004; Chen et al., 2005). Therefore, we suspect that SMT15 acts as a chloroplastic sulfate exporter to maintain sulfate homeostasis in Chlamydomonas cells. Under this assumption, we propose a model (Figure 9). In this model, defect in *SMT15* increases chloroplast sulfate concentration that subsequently enhances cysteine biosynthesis and facilitates glutathione biosynthesis. Even though it is not clear how cytosolic glutathione is reduced at S/M phase, increased influx of cytosolic glutathione in the *smt15-1* mutant may contribute to glutathione accumulation in the basal bodies. It is important to note that CMFDA-associated glutathione signal was also detected in the basal bodies of wt cells at S/M phase despite its low frequency (Table 4, Table 5, and Supplemental Table S1). It is therefore likely that glutathione in the basal bodies has a regulatory role during S/M phase in wt cells. This model needs to be further tested. Together, our studies reinforce the notion of the growing importance of glutathione-mediated redox balance in cell cycle regulation, and our data linking glutathione to cell division control is potentially of general significance that may apply to the animal and plant systems.

**Figure 9.**
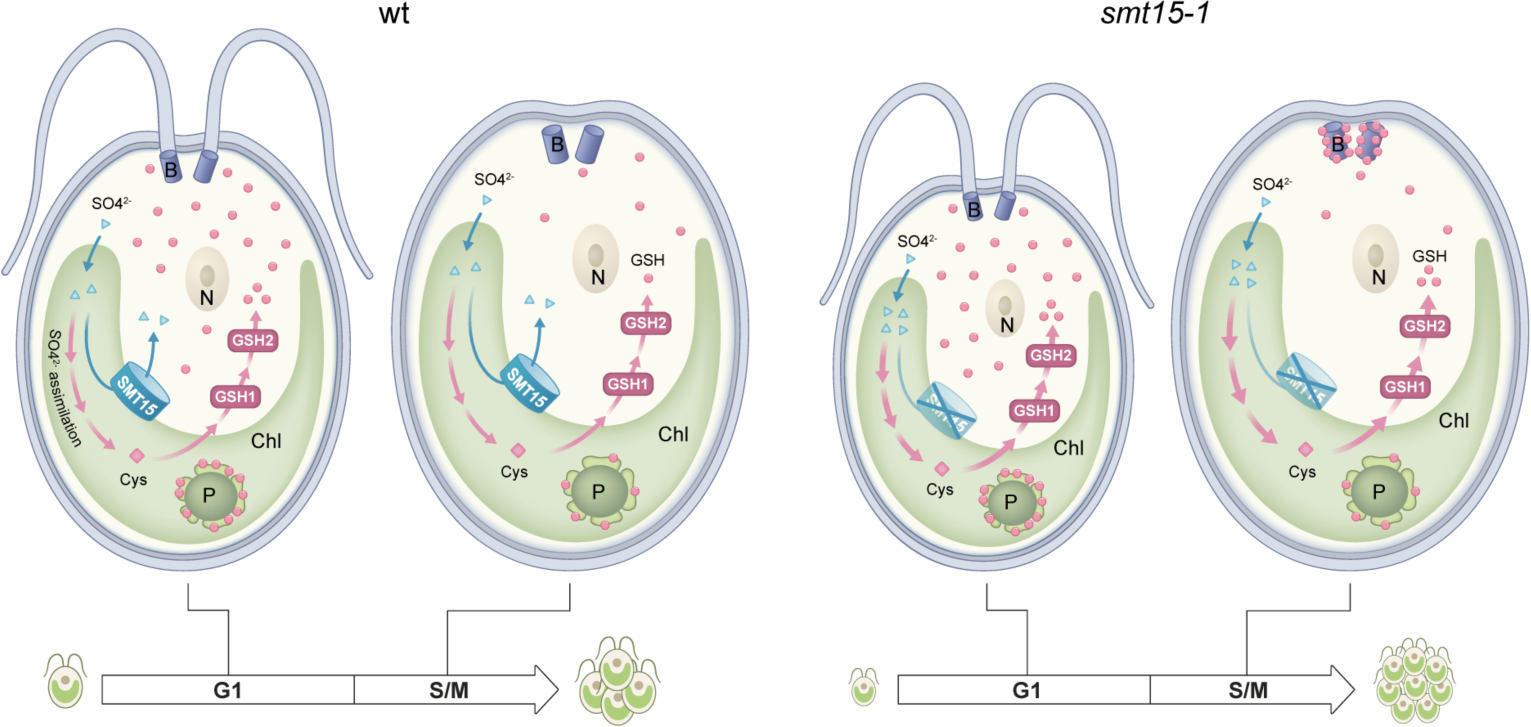
A model for SMT15-mediated glutathione regulation in Chlamydomonas. Lack of SMT15 on the chloroplast leads increased chloroplastic sulfate and facilitate glutathione biosynthesis in the cytosol. Increased flux of glutathione into the basal bodies at S/M phase increases the number of cell divisions and generates small-sized daughters. GSH1, γ-glutamylcysteine synthetase. GSH2, glutathione synthetase. Pink dot, glutathione (GSH). Blue triangle, SO4^2-^. Cys, cysteine. Chl, chloroplast. B, basal body. N, nucleus. P, pyrenoid.

## Materials and Methods

### Strains and growth conditions

*Chlamydomonas reinhardtii* strains 21gr (CC1690, MT^+^) and *smt15-1* (Fang and Umen, 2008; Fang et al., 2014) were used in the majority of experiments. Transgenic lines in this study were derived from 21gr, *smt15-1*, or *uvm4* (Neupert et al., 2009; Neupert et al., 2020) background.

Cells were grown in high salt medium (HSM, (Sueoka, 1960)) or Tris-acetate-phosphate (TAP, (Gorman and Levine, 1965)) under illumination at 250 to 300 μmol photons m^−2^ s^−1^ aerated with 0.5 % CO_2_. Culture synchronization was induced by growth in 14-h-light/10-h-dark cycles or 12-h-light/12-h-dark cycles as described.

### Strain Genotyping

To prepare crude genomic DNA for PCR, 1/3 loop-full of freshly-grown (3 to 5 days) cells were scraped from plates, resuspended in 50 µl TE buffer, and then incubated at ∼ 100 °C for 10 min. After centrifugation at highest speed in a microcentrifuge, one microliter of supernatant was used for PCR amplification. PCR fragments were amplified using Power Taq DNA polymerase (Genomics, Taiwan) in a final volume of 20 µl in the presence of 1X Power Taq buffer, 1 μM primers, 80 μM dNTP, 0.5 M Betaine, and 3% DMSO. Primer pairs used for PCR-based genotyping are listed in Table S2. PCR conditions were as follows: 96°C for 2 min, 42 cycles at 94°C for 30 s, 65°C for 30 s, and 72°C for 45s.

### Sulfate uptake assay

The DNA fragment encoding the six Histidine (6XHIS) tag with a polylinker (5’-AATTCTCTAGAGAGCTCGTCGACTTCACCACCACCACCACCACTAAAAGCTTCTCGA-3’) was ligated to the EcoRI and SalI digested pYX222x plasmid (Shibagaki et al., 2002). EcoRI and SalI digested *AtSULTR1;1* cDNA fragment (Shibagaki and Grossman, 2004) was then inserted into this modified pYX222x plasmid to allow translational fusion to the 6XHIS tag at the carboxyl-terminus. *AtSULTR1;1-6XHIS* cDNA was then removed by EcoRI and HindIII and cloned into pET28a vector to generate pET28a-AtSULTR1;1-6XHIS plasmid. For *SMT15*, the SacI and HindIII digested codon-optimized *SMT15*^316-1441^ cDNA was cloned into the modified pYX222x plasmid to allow translational fusion to the 6XHIS tag at the carboxyl-terminus. The *SMT15*^316-1441^*-6XHIS* cDNA was then digested by SacI and HindIII and cloned into pET28a vector to generate pET28a-SMT15^316-1441^-6XHIS plasmid.

For protein induction, 15 ng of pET28a, pET28a-AtSULTR1;1-6XHIS or pET28a-SMT15^316-1441^-6XHIS plasmid was transformed into *E. coli* Rosetta (DE3) cells by electroporation (Bio-Rad 1 mm cuvette, 18,000 V/cm, 200V, 25 mF). Transformed *E. coli* cells were selected on an LB agar plate containing 50 μg/mL kanamycin. A single colony was used to inoculate 200 ml LB broth containing 50 μg/mL kanamycin and incubated at 37°C until OD_600_ reached to approximately 0.5. For protein induction, 0.2 mM isopropyl β-D-1-thiogalactopyranoside (IPTG) was added to the culture and the culture was incubated at 25°C for an additional 3 h with vigorous shaking. Immunoblotting using the anti-polyHistidine (1:6000, GenScript A00186) was used to confirm protein expression.

^35^S-sulfate uptake was measured as described previously (Zolotarev et al., 2008) with slight modification. Briefly, IPTG-induced (3 h) *E. coli* cells were harvested in pre-weighed 50 ml tubes and washed once with 20 ml uptake medium (7 g/L K_2_HPO4, 3 g/L KH_2_PO4, 0.5g/L Na-Citrate, 0.12 g/L MgCl_2_·6H2O, 2.5 g/L NH_4_Cl, and 2 g/L glucose, pH 7.0). Washed cell pellets were resuspended in the uptake medium supplemented with 0.5 mM Na_2_SO_4_ and incubated at room temperature (RT) for 30 min, pelleted, and resuspended in equal amount of the uptake medium containing 0.1 mM Na_2_SO_4_. Twelve μCi of ^35^S-sulfate (Perkin Elmer) was added into 500 μl cell suspension by vortexing. One hundred microliter samples were removed at 0 and 10 min, and rapidly filtered through Millipore GSTF02500 filters mounted on a manifold. The filtered samples were then washed with 2.5 ml sulfate-free uptake medium twice and allowed to dry. ^35^S-sulfate was measured by liquid Scintillation Analyzer Tri-Carb 2810TR (Perkin Elmer).

### Generation of constructs and transgenic lines

The pF1 overexpression vector was modified from pSI103 (Sizova et al., 2001) and pEZ-GFP (Rasi et al., 2009) plasmids. Briefly, Primers 5′-ATTGCGGCCGCTCTAGACGGCGGGGA-3′ (carrying a NotI site) and 5′-ATTTCTAGACGCTTCAAATACGCCCAGCC-3′ (carrying a XbaI site) were used to amplify *aphVIII* cassette. NotI and XbaI digested *aphVIII* DNA fragment was inserted into pBluescript SK^−^. Primers 5′-AACTCTAGACACACACCTGCCCGTCTGCC-3′ (carrying a XbaI site) and 5′-AACGGTACCCACAGTCACGCTGTCTCCCC-3′ (carrying a KpnI site) were used to amplify PsaD expression cassette from pEZ plasmid (Rasi et al., 2009). XbaI and KpnI digested PsaD expression cassette was then inserted into pBluescript-aphVIII plasmid digested by XbaI and KpnI. The resulting plasmid is referred to as pF1. The eYFP cDNA fragment was PCR amplified and cloned into NdeI and EcoRI digested pF1 plasmid to make pF1-eYFP plasmid (Supplemental Figure S6).

For the SMT15-eYFP reporter constructs, the SMT15 N-terminal sequences were amplified by primers (Table S1) 88C12-C47NdeI and SMT15r1_BglII (20 amino acids), 88C12-C47NdeI and SMT15r2_BglII (46 amino acids), and 88C12-C47NdeI and SMT15r3_BglII (63 amino acids), digested by NdeI and BglII, and cloned into the pF1-eYFP plasmid. The constructs were validated by DNA sequencing. The ScaI-linearized construct was transformed into wild-type 21gr strain by electroporation as described previously (Kindle, 1990), and transformants were selected on TAP agar plates containing 20 μg/mL paromomycin. To screen for transgenic lines with eYFP expression, the paromomycin-resistant colonies were allowed to grow in TAP medium in a 96-well microtiter plate format at RT under continuous light for two days. The 96-well microtiter plate was then scanned for eYFP signal by SpectraMax iD3 Multi-Mode Microplate Reader (Molecular Devices). The eYFP positive transgenic lines were then validated by confocal microscope and immunoblotting analyses.

To generate pF1-GSH1-3XFLAG and pF1-GSH2-3XFLAG constructs, the full-length *GSH1* cDNA (Cre02.g077100) and *GSH2* cDNA (Cre17.g708800) were PCR amplified using 5′-TGAGCATATGGCTCTCGCCTCAGGCGTT-3′ and 5′-CTGCGAATTCGTACATGAACTCCTTGTACAGCGG-3′ (for *GSH1*), and 5′-GGCTCATATGCAGCACGAAATTCCTCGAG-3′ and 5′-AATCGAATTCCTCCACCAGGTAAGGGCTAT-3′ (for *GSH2*) from Chlamydomonas cDNA. The NdeI and EcoRI digested *GSH1* or *GSH2* cDNA was ligated into a NdeI/EcoRI digested pF1 plasmid to make pF1-GSH1-nsc and pF1-GSH2-nsc plasmids respectively. A 3XFLAG sequence was assembled from 5′-GCTAGCGACTACAAGGACCACGACGGCGACTACAAGGACCACGACATCGACTACA AGGAC-3′ and 5′-GCTAGCCTTGTCGTCGTCGTCCTTGTAGTCGATGTCGTGGTCCTTGTAGTCGCCGTC GTG-3’ oligos and PCR amplified using primers 5′-AATCGAATTCGCTAGCGACTACAAGGACCA-3′ and 5′-AATCCAATTGTCAGCTAGCCTTGTCGTCGT-3′. The EcoRI and MfeI digested 3XFLAG DNA fragment was ligated into the unique EcoRI site at the 3′ end of the *GSH1* or *GSH2* cDNA in pF1-GSH1-nsc or pF1-GSH2-nsc plasmid. The constructs, pF1-GSH1-3XFLAG and pF1-GSH2-3XFLAG, were sequenced to verify their accuracy. ScaI or ScaI/NotI digested pF1-GSH1-3XFLAG or pF1-GSH2-3XFLAG was transformed into 21gr strain by electroporation and the transformants were selected on paromomycin (20 μg/mL) containing plates.

The pF1-GSH1-lin-3XFLAG and pF1-GSH2-lin-3XFLAG constructs were modified from pF1-GSH1-nsc and pF1-GSH2-nsc constructs respectively. Briefly, the synthesized linker and 3XFLAG DNA sequence was ligated into EcoRI digested pF1-GSH1 or pF1-GSH2 to make pF1-GSH1-lin-3XFLAG and pF1-GSH2-lin-3XFLAG constructs respectively. Both pF1-GSH1-lin-3XFLAG and pF1-GSH2-lin-3XFLAG constructs were sequenced to verify their accuracy. KpnI digested pF1-GSH1-lin-3XFLAG or pF1-GSH2-lin-3XFLAG was transformed into *uvw4* strain (Neupert et al., 2009) by glass bead transformation (Kindle, 1990) and the transformants were selected on paromomycin (20 μg/mL) containing plates.

### Immunoblotting

Approximately 3 × 10^7^ cells were collected by centrifugation at 1181 × g for 5 min at RT. Cells were resuspended in 200 µL of 1× Urea protein extraction buffer (250 mM Tris-HCl [pH 6.8], 3.5% [w/v] SDS, 10% [v/v] Glycerol, 1M Urea) supplemented with 1× protease inhibitor cocktail (Sigma-Aldrich, P9599), 1 mM PMSF, 1 mM benzamidine and 10 µM MG132 and incubated on ice for 15 min. After centrifugation at 16200 × g for 30 min at 4°C, the supernatant protein extract was collected and protein concentration was determined using DC Protein Assay Kit (Bio-Rad).

Approximately 100 µg protein of each sample was separated by 12% SDS-PAGE and transferred to an Immobilon-PVDF membrane (Merck Millipore). The blot was blocked with 5% non-fat milk in 1× TBST (20 mM Tris base, 150 mM NaCl, 0.1% [v/v] Tween 20) for 1 h at RT. The blot was the incubated with GFP antibody (1:1000, Roche) diluted in 5% non-fat milk at 4°C overnight. The following day, the blot was washed 3 times with 1× TBST for 15 min each at RT and then incubated with horseradish peroxidase–conjugated goat anti-mouse IgG antibody (1:20000, Jackson ImmunoResearch 115-035-003) diluted in 5% non-fat milk for 1 h at RT. After incubation, the blot was washed 3 times with 1× TBST for 15 min each at RT. For chemiluminescence detection, the luminol Advansta WesternBright ECL was applied to the blot and images were captured by ChemiDox XRS+ Imager (Bio-Rad).

### Glutathione measurements

The total glutathione content was measured using a Glutathione Assay Kit (Cayman Chemical) following the manufacturer’s instructions. Briefly, ∼5 × 10^7^ cells were harvested and washed with 10 ml of ice-cold 1× PBS (137 mM NaCl, 2.7 mM KCl, 10 mM Na_2_HPO_4_, 1.8 mM KH_2_PO_4_) twice. For measurement of total glutathione, the cell pellets were resuspended in 500 μl of ice-cold 5% [w/v] 5-sulfo-salicylic acid dehydrate (SSA) and homogenized by vortexing in the presence of 300 µL zirconium beads (BioSpec Products) at 4°C for 30 min. The cell lysate was centrifuged at 21,130 × g for 5 min at 4°C to remove cell debris. The supernatant was snap frozen in liquid N_2_ and stored at −75°C before analysis. Glutathione reductase was added to reduce GSSG to GSH and total GSH was measured as described in the instruction manual. GSH content was extrapolated based on a standard curve generated by GSSG with a known concentration. Total glutathione was normalized to total protein content. For protein content measurement, ∼1 × 10^7^ cells were collected and boiled in 600 µL of 2% [w/v] SDS solution at 100°C for 10 min. Detergent-compatible (DC) protein assay kit (Bio-Rad) was used to determine protein content.

### Cell size and cell cycle analysis

Cell size was measured directly by a Coulter Counter (Multisizer 3; Beckman-Coulter, Miami, Florida, United States). Mitotic index assay was determined as described previously (Umen and Goodenough, 2001). For cell cycle analysis, cells were fixed as described previously (Fang et al., 2006). Briefly, 10 ml of cells was pelleted in the presence of 0.005% Tween 20 and fixed by resuspending in 10 ml of ethanol/acetic acid (3:1) at RT for 1 h. Fixed cells were then washed once with 10 ml of FACS buffer (0.2 M Tris [pH 7.5], 20 mM EDTA), resuspended in 1 ml of FACS buffer, and stored at 4°C. Prior to image cytometry, cells were incubated in FACS buffer at 37°C with 100 μg/ml RNase A for 2 h, washed once with 1 ml of PBS, and resuspended in 1 ml of PBS. DNA was stained with 0.5 nM Sytox Green (Invitrogen), and cytometry was performed with a NucleoCounter NC-3000 system and analyzed using Modfit LT software (Chemometec). For the cell division behavior analysis presented in Figure 5F, 14 h light-grown cells were plated on HSM agar plates and allowed to incubate in the dark. Cell division numbers for individual dark-incubated cells were measured (Fang et al., 2006). At least 220 cells were measured for each strain.

### RT-qPCR

Total RNA was isolated as previously described (Fang et al., 2014). Five micrograms of RNA was used for cDNA synthesis. cDNAs were reverse transcribed in the presence of a mixture of oligo dT and random primers (9:1) at 50°C for 50 min using ThermoScript Reverse Transcriptase or SuperScript IV Reverse Transcriptase (Invitrogen) following the manufacturer’s instructions. Each 20 mL PCR reaction contained 1 mL of 1/20-fold diluted cDNA, 1X Phusion GC buffer, 0.2 mM deoxynucleotide triphosphate (dNTP), 1 mM primers, 3% (v/v) DMSO, 0.5 M betaine, and 0.4 units Phusion High-Fidelity DNA polymerase (New England Biolabs). The qPCR program was performed as follows: 98°C for 1min, 40 cycles of 98°C for 5s, 63°C for 20s. The primers used for PCR are listed in Table S3.

### 5-chloromethylfluorescein diacetate (CMFDA) staining

Approximately 1 × 10^7^ cells were harvested by centrifugation at 1000 × g for 5 min at RT. The cell pellet was resuspended in 1 mL HSM. Half a microliter of 10 mM CMFDA or 0.5 µL DMSO (as the negative control) was added to 1 mL cell suspension. The cells were mixed by rotating in the dark at RT for 45 min. The cells were then harvested by centrifugation at 1000 × g for 5 min at RT. The cell pellets were washed twice with 1 ml of 1 × PBS. A Zeiss 980 confocal microscope with the following excitation and emission settings was used to capture the fluorescent signals: CMFDA, 488 nm excitation with 500-544 nm emission; chlorophyll, 548 nm excitation with 640-700 nm emission.

### Immunofluorescence analysis

Approximately 1 × 10^7^ cells were harvested by centrifugation at 1181 × g for 5 min at RT. The cover slips were treated with poly-L-lysin solution (Sigma-Aldrich P8920) at RT for 30 min. Cells were allowed to attach to poly-L-lysin-coated cover slips at RT for 15 min. The attached cells were fixed by immersion into two changes of pre-cooled −20°C 200 μL 100% (v/v) methanol for 10 min and rehydrated in 500 µL 1× PBS (137 mM NaCl, 2.7 mM KCl, 10 mM Na_2_HPO_4_, 1.8 mM KH_2_PO_4_) at RT for 15 min. Cover slips were then blocked in 150 μL of blocking solution (5% [w/v] BSA, 5% [v/v] Normal Goat Serum [Jackson ImmunoResearch Laboratories] in 1× PBS) at RT for 90 min. After blocking, the cover slips were incubated with 150 μL of primary antibody (1:50 anti-FLAG M2 antibody, Sigma-Aldrich F1804) diluted in blocking solution overnight at 4°C. Next day, the cover slips were washed 3 times with 1× PBST (1× PBS supplemented with 0.1% [v/v] Tween 20) at RT for 10 min. The cover slips were then incubated with 150 µL 1:500 secondary antibody (Alexa Fluor 488-conjugated Goat anti-mouse IgG, Thermo Fisher Scientific) diluted in blocking solution in the dark at RT for 120 min. After incubation, the cover slips were washed with 1× PBS once. To stain DNA, the cover slips were incubated with 150 µL 4′,6-diamidino-2-phenylindole (DAPI, 2µg/ml) solution diluted in 1× PBS in the dark at RT for 10 min. Following four washes with 1× PBST for 10 min per wash at RT, cover slips were mounted with Vectashield Antifade Mounting Medium (vector laboratories H-1000) to prevent photobleaching. A Zeiss 980 confocal microscope with filter sets (488 nm laser for Alexa Fluor 488; 405 nm UV light laser for DAPI) were used to capture the images.

## Acknowledgements

We thank the Confocal Microscopy at Academia Sinica Biotechnology Center in Southern Taiwan for the service, Mr. Chin-Lin Chung for technical support, Ms. Miranda Loney for English editing, and Dr. Ying Wang for graphic design of Figure 9.

## Author contributions

SCF conceived the idea and coordinated the study. SCF and PJH designed the experiments and wrote the manuscript. PJH, CHC, YLL and HYL conducted experiments and analyzed the data. All the authors have read and approved the final manuscript.

## References

Ahn E, Kumar P, Mukha D, Tzur A, Shlomi T (2017) Temporal fluxomics reveals oscillations in TCA cycle flux throughout the mammalian cell cycle. Mol Syst Biol 13: 953

Ball L, Accotto GP, Bechtold U, Creissen G, Funck D, Jimenez A, Kular B, Leyland N, Mejia-Carranza J, Reynolds H, Karpinski S, Mullineaux PM (2004) Evidence for a direct link between glutathione biosynthesis and stress defense gene expression in Arabidopsis. Plant Cell 16: 2448–2462

Blackburn K, Bustamante-Marin X, Yin W, Goshe MB, Ostrowski LE (2017) Quantitative proteomic analysis of human airway cilia identifies previously uncharacterized proteins of high abundance. J Proteome Res 16: 1579–1592

Boonstra J, Post JA (2004) Molecular events associated with reactive oxygen species and cell cycle progression in mammalian cells. Gene 337: 1–13

Burch PM, Heintz NH (2005) Redox regulation of cell-cycle re-entry: cyclin D1 as a primary target for the mitogenic effects of reactive oxygen and nitrogen species. Antioxid Redox Signal 7: 741–751

Burhans WC, Heintz NH (2009) The cell cycle is a redox cycle: linking phase-specific targets to cell fate. Free Radic Biol Med 47: 1282–1293

Cairns NG, Pasternak M, Wachter A, Cobbett CS, Meyer AJ (2006) Maturation of Arabidopsis seeds is dependent on glutathione biosynthesis within the embryo. Plant Physiol 141: 446–455

Chen H-C, Melis A (2004) Localization and function of SulP, a nuclear-encoded chloroplast sulfate permease in *Chlamydomonas reinhardtii*. Planta 220: 198–210

Chen HC, Newton AJ, Melis A (2005) Role of SulP, a nuclear-encoded chloroplast sulfate permease, in sulfate transport and H2 evolution in *Chlamydomonas reinhardtii*. Photosynth Res 84: 289–296

Chen HC, Yokthongwattana K, Newton AJ, Melis A (2003) SulP, a nuclear gene encoding a putative chloroplast-targeted sulfate permease in *Chlamydomonas reinhardtii*. Planta 218: 98–106

Chen J, Yang L, Yan X, Liu Y, Wang R, Fan T, Ren Y, Tang X, Xiao F, Liu Y, Cao S (2016) Zinc-finger transcription factor ZAT6 positively regulates cadmium tolerance through the glutathione-dependent pathway in Arabidopsis. Plant Physiol 171: 707–719

Chen Z, Odstrcil EA, Tu BP, McKnight SL (2007) Restriction of DNA replication to the reductive phase of the metabolic cycle protects genome integrity. Science 316: 1916–1919

Chiu J, Dawes IW (2012) Redox control of cell proliferation. Trends Cell Biol 22: 592–601

Cobbett CS, May MJ, Howden R, Rolls B (1998) The glutathione-deficient, cadmium-sensitive mutant, *cad2-1*, of *Arabidopsis thaliana* is deficient in gamma-glutamylcysteine synthetase. Plant J 16: 73–78

Craigie RA, Cavaliersmith T (1982) Cell volume and the control of the *Chlamydomonas* cell cycle. J Cell Sci 54: 173–191

Creissen G, Firmin J, Fryer M, Kular B, Leyland N, Reynolds H, Pastori G, Wellburn F, Baker N, Wellburn A, Mullineaux P (1999) Elevated glutathione biosynthetic capacity in the chloroplasts of transgenic tobacco plants paradoxically causes increased oxidative stress. Plant Cell 11: 1277–1292

de Simone A, Hubbard R, Vinegra de la Torre N, Velappan Y, Wilson M, Considine MJ, Soppe WJJ, Foyer CH (2017) Redox changes during the cell cycle in the embryonic root meristem of *Arabidopsis thaliana*. Antioxid Redox Signal 27: 1505–1519

Diaz Vivancos P, Dong Y, Ziegler K, Markovic J, Pallardó FV, Pellny TK, Verrier PJ, Foyer CH (2010) Recruitment of glutathione into the nucleus during cell proliferation adjusts whole-cell redox homeostasis in *Arabidopsis thaliana* and lowers the oxidative defence shield. Plant J 64: 825–838

Diaz Vivancos P, Wolff T, Markovic J, Pallardó FV, Foyer CH (2010) A nuclear glutathione cycle within the cell cycle. Biochem J 431: 169–178

Diaz-Vivancos P, de Simone A, Kiddle G, Foyer CH (2015) Glutathione--linking cell proliferation to oxidative stress. Free Radic Biol Med 89: 1154–1164

Dolzblasz A, Smakowska E, Gola EM, Sokolowska K, Kicia M, Janska H (2016) The mitochondrial protease AtFTSH4 safeguards Arabidopsis shoot apical meristem function. Sci Rep 6: 28315

Donnan L, John PC (1983) Cell cycle control by timer and sizer in *Chlamydomonas*. Nature 304: 630–633

Dutcher SK (2003) Elucidation of basal body and centriole functions in *Chlamydomonas reinhardtii*. Traffic 4: 443–451

Dutcher SK, Trabuco EC (1998) The UNI3 gene is required for assembly of basal bodies of Chlamydomonas and encodes delta-tubulin, a new member of the tubulin superfamily. Mol Biol Cell 9: 1293–1308

Ehler LL, Holmes JA, Dutcher SK (1995) Loss of spatial control of the mitotic spindle apparatus in a *Chlamydomonas reinhardtii* mutant strain lacking basal bodies. Genetics 141: 945–960

Fang SC, Chung CL, Chen CH, Lopez-Paz C, Umen JG (2014) Defects in a new class of sulfate/anion transporter link sulfur acclimation responses to intracellular glutathione levels and cell cycle control. Plant Physiol 166: 1852–1868

Fang SC, de los Reyes C, Umen JG (2006) Cell size checkpoint control by the retinoblastoma tumor suppressor pathway. PLoS Genet 2: e167

Fang SC, Umen JG (2008) A suppressor screen in Chlamydomonas identifies novel components of the retinoblastoma tumor suppressor pathway. Genetics 178: 1295–1310

García-Giménez JL, Markovic J, Dasi F, Queval G, Schnaubelt D, Foyer CH, Pallardó FV (2013) Nuclear glutathione. Biochim Biophys Acta 1830: 3304–3316

Gorman DS, Levine RP (1965) Cytochrome f and plastocyanin: their sequence in the photosynthetic electron transport chain of *Chlamydomonas reinhardii*. Proc Natl Acad Sci U S A 54: 1665–1669

Goyal U, Renvoise B, Chang J, Blackstone C (2014) Spastin-interacting protein NA14/SSNA1 functions in cytokinesis and axon development. PLoS One 9: e112428

Griffith OW, Meister A (1985) Origin and turnover of mitochondrial glutathione. Proc Natl Acad Sci U S A 82: 4668–4672

Hartl J, Kiefer P, Kaczmarczyk A, Mittelviefhaus M, Meyer F, Vonderach T, Hattendorf B, Jenal U, Vorholt JA (2020) Untargeted metabolomics links glutathione to bacterial cell cycle progression. Nat Metab 2: 153–166

Hoffman A, Spetner LM, Burke M (2008) Ramifications of a redox switch within a normal cell: its absence in a cancer cell. Free Radic Biol Med 45: 265–268

Ivanova JS, Pugovkina NA, Neganova IE, Kozhukharova IV, Nikolsky NN, Lyublinskaya OG (2021) Cell cycle-coupled changes in the level of reactive oxygen species support the proliferation of human pluripotent stem cells. Stem Cells 39: 1671–1687

Jackson PK (2011) Do cilia put brakes on the cell cycle? Nat Cell Biol 13: 340–342

Ji Y, Chae S, Lee HK, Park I, Kim C, Ismail T, Kim Y, Park JW, Kwon OS, Kang BS, Lee DS, Bae JS, Kim SH, Moon PG, Baek MC, Park MJ, Kil IS, Rhee SG, Kim J, Huh YH, Shin JY, Min KJ, Kwon TK, Jang DG, Woo HA, Kwon T, Park TJ, Lee HS (2019) Peroxiredoxin5 controls vertebrate ciliogenesis by modulating mitochondrial reactive oxygen species. Antioxid Redox Signal 30: 1731–1745

Jiang X, Yu Y, Chen J, Zhao M, Chen H, Song X, Matzuk AJ, Carroll SL, Tan X, Sizovs A, Cheng N, Wang MC, Wang J (2015) Quantitative imaging of glutathione in live cells using a reversible reaction-based ratiometric fluorescent probe. ACS Chem Biol 10: 864–874

Jiao GZ, Cao XY, Cui W, Lian HY, Miao YL, Wu XF, Han D, Tan JH (2013) Developmental potential of prepubertal mouse oocytes is compromised due mainly to their impaired synthesis of glutathione. PLoS One 8: e58018

Keller LC, Wemmer KA, Marshall WF (2010) Influence of centriole number on mitotic spindle length and symmetry. Cytoskeleton (Hoboken) 67: 504–518

Kim S, Zaghloul NA, Bubenshchikova E, Oh EC, Rankin S, Katsanis N, Obara T, Tsiokas L (2011) Nde1-mediated inhibition of ciliogenesis affects cell cycle re-entry. Nat Cell Biol 13: 351–360

Kindle KL (1990) High-frequency nuclear transformation of *Chlamydomonas reinhardtii*. Proc Natl Acad Sci USA 87: 1228–1232

Kirova DG, Judasova K, Vorhauser J, Zerjatke T, Leung JK, Glauche I, Mansfeld J (2022) A ROS-dependent mechanism promotes CDK2 phosphorylation to drive progression through S phase. Dev Cell 57: 1712–1727 e1719

Lantz RC, Lemus R, Lange RW, Karol MH (2001) Rapid reduction of intracellular glutathione in human bronchial epithelial cells exposed to occupational levels of toluene diisocyanate. Toxicol Sci 60: 348–355

Lee SR, Kwon KS, Kim SR, Rhee SG (1998) Reversible inactivation of protein-tyrosine phosphatase 1B in A431 cells stimulated with epidermal growth factor. J Biol Chem 273: 15366–15372

Li A, Saito M, Chuang JZ, Tseng YY, Dedesma C, Tomizawa K, Kaitsuka T, Sung CH (2011) Ciliary transition zone activation of phosphorylated Tctex-1 controls ciliary resorption, S-phase entry and fate of neural progenitors. Nat Cell Biol 13: 402–411

Lim DC, Joukov V, Rettenmaier TJ, Kumagai A, Dunphy WG, Wells JA, Yaffe MB (2020) Redox priming promotes Aurora A activation during mitosis. Sci Signal 13

Lim DC, Joukov V, Yaffe MB (2020) Are redox changes a critical switch for mitotic progression? Mol Cell Oncol 7: 1832419

Lim JM, Lee KS, Woo HA, Kang D, Rhee SG (2015) Control of the pericentrosomal H2O2 level by peroxiredoxin I is critical for mitotic progression. J Cell Biol 210: 23–33

Lin YL, Chung CL, Chen MH, Chen CH, Fang SC (2020) SUMO protease SMT7 modulates ribosomal protein L30 and regulates cell-size checkpoint function. Plant Cell 32: 1285–1307

Lyublinskaya OG, Borisov YG, Pugovkina NA, Smirnova IS, Obidina JV, Ivanova JS, Zenin VV, Shatrova AN, Borodkina AV, Aksenov ND, Zemelko VI, Burova EB, Puzanov MV, Nikolsky NN (2015) Reactive oxygen species are required for human mesenchymal stem cells to initiate proliferation after the quiescence exit. Oxid Med Cell Longev 2015: 502105

Mackinder LC, Meyer MT, Mettler-Altmann T, Chen VK, Mitchell MC, Caspari O, Freeman Rosenzweig ES, Pallesen L, Reeves G, Itakura A, Roth R, Sommer F, Geimer S, Muhlhaus T, Schroda M, Goodenough U, Stitt M, Griffiths H, Jonikas MC (2016) A repeat protein links Rubisco to form the eukaryotic carbon-concentrating organelle. Proc Natl Acad Sci U S A 113: 5958–5963

Macleod KF (2008) The role of the RB tumour suppressor pathway in oxidative stress responses in the haematopoietic system. Nat Rev Cancer 8: 769–781

Mandigo AC, Yuan W, Xu K, Gallagher P, Pang A, Guan YF, Shafi AA, Thangavel C, Sheehan B, Bogdan D, Paschalis A, McCann JJ, Laufer TS, Gordon N, Vasilevskaya IA, Dylgjeri E, Chand SN, Schiewer MJ, Domingo-Domenech J, Den RB, Holst J, McCue PA, de Bono JS, McNair C, Knudsen KE (2021) RB/E2F1 as a master regulator of cancer cell metabolism in advanced disease. Cancer Discov 11: 2334–2353

Markovic J, Borrás C, Ortega A, Sastre J, Viña J, Pallardó FV (2007) Glutathione is recruited into the nucleus in early phases of cell proliferation. J Biol Chem 282: 20416–20424

Markovic J, Mora NJ, Broseta AM, Gimeno A, de-la-Concepción N, Viña J, Pallardó FV (2009) The depletion of nuclear glutathione impairs cell proliferation in 3t3 fibroblasts. PLoS ONE 4: e6413

Martindale JL, Holbrook NJ (2002) Cellular response to oxidative stress: signaling for suicide and survival. J Cell Physiol 192: 1–15

Mata M, Sarrion I, Armengot M, Carda C, Martinez I, Melero JA, Cortijo J (2012) Respiratory syncytial virus inhibits ciliagenesis in differentiated normal human bronchial epithelial cells: effectiveness of N-acetylcysteine. PLoS One 7: e48037

Matsuura K, Lefebvre PA, Kamiya R, Hirono M (2004) Bld10p, a novel protein essential for basal body assembly in Chlamydomonas: localization to the cartwheel, the first ninefold symmetrical structure appearing during assembly. J Cell Biol 165: 663–671

Menon SG, Goswami PC (2007) A redox cycle within the cell cycle: ring in the old with the new. Oncogene 26: 1101–1109

Merchant SS, Prochnik SE, Vallon O, Harris EH, Karpowicz SJ, Witman GB, Terry A, Salamov A, Fritz-Laylin LK, Marechal-Drouard L, Marshall WF, Qu LH, Nelson DR, Sanderfoot AA, Spalding MH, Kapitonov VV, Ren Q, Ferris P, Lindquist E, Shapiro H, Lucas SM, Grimwood J, Schmutz J, Cardol P, Cerutti H, Chanfreau G, Chen CL, Cognat V, Croft MT, Dent R, Dutcher S, Fernandez E, Fukuzawa H, González-Ballester D, González-Halphen D, Hallmann A, Hanikenne M, Hippler M, Inwood W, Jabbari K, Kalanon M, Kuras R, Lefebvre PA, Lemaire SD, Lobanov AV, Lohr M, Manuell A, Meier I, Mets L, Mittag M, Mittelmeier T, Moroney JV, Moseley J, Napoli C, Nedelcu AM, Niyogi K, Novoselov SV, Paulsen IT, Pazour G, Purton S, Ral JP, Riano-Pachon DM, Riekhof W, Rymarquis L, Schroda M, Stern D, Umen J, Willows R, Wilson N, Zimmer SL, Allmer J, Balk J, Bisova K, Chen CJ, Elias M, Gendler K, Hauser C, Lamb MR, Ledford H, Long JC, Minagawa J, Page MD, Pan J, Pootakham W, Roje S, Rose A, Stahlberg E, Terauchi AM, Yang P, Ball S, Bowler C, Dieckmann CL, Gladyshev VN, Green P, Jorgensen R, Mayfield S, Mueller-Roeber B, Rajamani S, Sayre RT, Brokstein P, Dubchak I, Goodstein D, Hornick L, Huang YW, Jhaveri J, Luo Y, Martinez D, Ngau WC, Otillar B, Poliakov A, Porter A, Szajkowski L, Werner G, Zhou K, Grigoriev IV, Rokhsar DS, Grossman AR (2007) The *Chlamydomonas* genome reveals the evolution of key animal and plant functions. Science 318: 245–250

Mhamdi A, Hager J, Chaouch S, Queval G, Han Y, Taconnat L, Saindrenan P, Gouia H, Issakidis-Bourguet E, Renou JP, Noctor G (2010) Arabidopsis GLUTATHIONE REDUCTASE1 plays a crucial role in leaf responses to intracellular hydrogen peroxide and in ensuring appropriate gene expression through both salicylic acid and jasmonic acid signaling pathways. Plant Physiol 153: 1144–1160

Neupert J, Gallaher SD, Lu Y, Strenkert D, Segal N, Barahimipour R, Fitz-Gibbon ST, Schroda M, Merchant SS, Bock R (2020) An epigenetic gene silencing pathway selectively acting on transgenic DNA in the green alga Chlamydomonas. Nat Commun 11: 6269

Neupert J, Karcher D, Bock R (2009) Generation of Chlamydomonas strains that efficiently express nuclear transgenes. Plant J 57: 1140–1150

Nicolay BN, Gameiro PA, Tschop K, Korenjak M, Heilmann AM, Asara JM, Stephanopoulos G, Iliopoulos O, Dyson NJ (2013) Loss of RBF1 changes glutamine catabolism. Genes Dev 27: 182–196

Noctor G, Arisi AC, Jouanin L, Foyer CH (1998) Manipulation of glutathione and amino acid biosynthesis in the chloroplast. Plant Physiol 118: 471–482

Noctor G, Strohm M, Jouanin L, Kunert KJ, Foyer CH, Rennenberg H (1996) Synthesis of glutathione in leaves of transgenic poplar overexpressing g-glutamylcysteine synthetase. Plant Physiol 112: 1071–1078

O’Toole ET, Dutcher SK (2014) Site-specific basal body duplication in Chlamydomonas. Cytoskeleton (Hoboken) 71: 108–118

Odom RY, Dansby MY, Rollins-Hairston AM, Jackson KM, Kirlin WG (2009) Phytochemical induction of cell cycle arrest by glutathione oxidation and reversal by N-acetylcysteine in human colon carcinoma cells. Nutr Cancer 61: 332–339

Olson BJ, Oberholzer M, Li Y, Zones JM, Kohli HS, Bisova K, Fang SC, Meisenhelder J, Hunter T, Umen JG (2010) Regulation of the *Chlamydomonas* cell cycle by a stable, chromatin-associated retinoblastoma tumor suppressor complex. Plant Cell 22: 3331–3347

Pallardó FV, Markovic J, García JL, Viña J (2009) Role of nuclear glutathione as a key regulator of cell proliferation. Mol Aspects Med 30: 77–85

Pan J, Snell WJ (2000) Regulated targeting of a protein kinase into an intact flagellum. An aurora/Ipl1p-like protein kinase translocates from the cell body into the flagella during gamete activation in chlamydomonas. J Biol Chem 275: 24106–24114

Pasternak M, Lim B, Wirtz M, Hell R, Cobbett CS, Meyer AJ (2008) Restricting glutathione biosynthesis to the cytosol is sufficient for normal plant development. Plant J 53: 999–1012

Patron NJ, Durnford DG, Kopriva S (2008) Sulfate assimilation in eukaryotes: fusions, relocations and lateral transfers. BMC Evolutionary Biology 8: 1–14

Patterson JC, Joughin BA, van de Kooij B, Lim DC, Lauffenburger DA, Yaffe MB (2019) ROS and oxidative stress are elevated in mitosis during asynchronous cell cycle progression and are exacerbated by mitotic arrest. Cell Syst 8: 163–167 e162

Pecani K, Lieberman K, Tajima-Shirasaki N, Onishi M, Cross FR (2022) Control of division in Chlamydomonas by cyclin B/CDKB1 and the anaphase-promoting complex. PLoS Genet 18: e1009997

Plummer JL, Smith BR, Sies H, Bend JR (1981) Chemical depletion of glutathione *in vivo*. Methods Enzymol 77: 50–59

Price ME, Sisson JH (2019) Redox regulation of motile cilia in airway disease. Redox Biol 27: 101146

Rasi MQ, Parker JD, Feldman JL, Marshall WF, Quarmby LM (2009) Katanin knockdown supports a role for microtubule severing in release of basal bodies before mitosis in Chlamydomonas. Mol Biol Cell 20: 379–388

Ravina CG, Chang CI, Tsakraklides GP, McDermott JP, Vega JM, Leustek T, Gotor C, Davies JP (2002) The *sac* mutants of *Chlamydomonas reinhardtii* reveal transcriptional and posttranscriptional control of cysteine biosynthesis. Plant Physiol 130: 2076–2084

Reddy Chichili VP, Kumar V, Sivaraman J (2013) Linkers in the structural biology of protein-protein interactions. Protein Sci 22: 153–167

Sabourin M, Tuzon CT, Fisher TS, Zakian VA (2007) A flexible protein linker improves the function of epitope-tagged proteins in *Saccharomyces cerevisiae*. Yeast 24: 39–45

Sarsour EH, Kumar MG, Chaudhuri L, Kalen AL, Goswami PC (2009) Redox control of the cell cycle in health and disease. Antioxid Redox Signal 11: 2985–3011

Shanmugam V, Tsednee M, Yeh KC (2012) *ZINC TOLERANCE INDUCED BY IRON 1* reveals the importance of glutathione in the cross-homeostasis between zinc and iron in *Arabidopsis thaliana*. Plant J 69: 1006–1017

Shanmugam V, Wang YW, Tsednee M, Karunakaran K, Yeh KC (2015) Glutathione plays an essential role in nitric oxide-mediated iron-deficiency signaling and iron-deficiency tolerance in Arabidopsis. Plant J 84: 464–477

Shibagaki N, Grossman AR (2004) Probing the function of STAS domains of the Arabidopsis sulfate transporters. J Biol Chem 279: 30791–30799

Shibagaki N, Rose A, McDermott JP, Fujiwara T, Hayashi H, Yoneyama T, Davies JP (2002) Selenate-resistant mutants of Arabidopsis thaliana identify Sultr1;2, a sulfate transporter required for efficient transport of sulfate into roots. Plant J 29: 475–486

Silflow CD, LaVoie M, Tam LW, Tousey S, Sanders M, Wu W, Borodovsky M, Lefebvre PA (2001) The Vfl1 Protein in Chlamydomonas localizes in a rotationally asymmetric pattern at the distal ends of the basal bodies. J Cell Biol 153: 63–74

Simet SM, Pavlik JA, Sisson JH (2013) Proteomic analysis of bovine axonemes exposed to acute alcohol: role of endothelial nitric oxide synthase and heat shock protein 90 in cilia stimulation. Alcohol Clin Exp Res 37: 609–615

Sizova I, Fuhrmann M, Hegemann P (2001) A *Streptomyces rimosus aphVIII* gene coding for a new type phosphotransferase provides stable antibiotic resistance to *Chlamydomonas reinhardtii*. Gene 277: 221–229

Sueoka N (1960) Mitotic replication of deoxyribonucleic acid in *Chlamydomonas reinhardii*. Proc Natl Acad Sci U S A 46: 83–91

Toledano MB, Delaunay-Moisan A, Outten CE, Igbaria A (2013) Functions and cellular compartmentation of the thioredoxin and glutathione pathways in yeast. Antioxid Redox Signal 18: 1699–1711

Trujillo-Hernandez JA, Bariat L, Enders TA, Strader LC, Reichheld JP, Belin C (2020) A glutathione-dependent control of the indole butyric acid pathway supports Arabidopsis root system adaptation to phosphate deprivation. J Exp Bot 71: 4843–4857

Tsukagoshi H, Busch W, Benfey PN (2010) Transcriptional regulation of ROS controls transition from proliferation to differentiation in the root. Cell 143: 606–616

Tu BP, Kudlicki A, Rowicka M, McKnight SL (2005) Logic of the yeast metabolic cycle: temporal compartmentalization of cellular processes. Science 310: 1152–1158

Umen JG (2005) The elusive sizer. Curr Opin Cell Biol 17: 435–441

Umen JG, Goodenough UW (2001) Control of cell division by a retinoblastoma protein homolog in *Chlamydomonas*. Genes Dev 15: 1652–1661

Vander Heiden MG, Cantley LC, Thompson CB (2009) Understanding the Warburg effect: the metabolic requirements of cell proliferation. Science 324: 1029–1033

Vernoux T, Wilson RC, Seeley KA, Reichheld JP, Muroy S, Brown S, Maughan SC, Cobbett CS, Van Montagu M, Inzé D, May MJ, Sung ZR (2000) The *ROOT MERISTEMLESS1/CADMIUM SENSITIVE2* gene defines a glutathione-dependent pathway involved in initiation and maintenance of cell division during postembryonic root development. The Plant cell 12: 97–110

Voehringer DW, McConkey DJ, McDonnell TJ, Brisbay S, Meyn RE (1998) Bcl-2 expression causes redistribution of glutathione to the nucleus. Proc Natl Acad Sci U S A 95: 2956–2960

Wachter A, Wolf S, Steininger H, Bogs J, Rausch T (2005) Differential targeting of GSH1 and GSH2 is achieved by multiple transcription initiation: implications for the compartmentation of glutathione biosynthesis in the Brassicaceae. Plant J 41: 15–30

Wang H, Gau B, Slade WO, Juergens M, Li P, Hicks LM (2014) The global phosphoproteome of *Chlamydomonas reinhardtii* reveals complex organellar phosphorylation in the flagella and thylakoid membrane. Mol Cell Proteomics 13: 2337–2353

Wang H, Nicolay BN, Chick JM, Gao X, Geng Y, Ren H, Gao H, Yang G, Williams JA, Suski JM, Keibler MA, Sicinska E, Gerdemann U, Haining WN, Roberts TM, Polyak K, Gygi SP, Dyson NJ, Sicinski P (2017) The metabolic function of cyclin D3-CDK6 kinase in cancer cell survival. Nature 546: 426–430

Willems E, Dedobbeleer M, Digregorio M, Lombard A, Lumapat PN, Rogister B (2018) The functional diversity of Aurora kinases: a comprehensive review. Cell Div 13: 7

Wu G, Fang YZ, Yang S, Lupton JR, Turner ND (2004) Glutathione metabolism and its implications for health. J Nutr 134: 489–492

Xue C, Botkin SJ, Johns RA (1996) Localization of endothelial NOS at the basal microtubule membrane in ciliated epithelium of rat lung. J Histochem Cytochem 44: 463–471

Yu Q, Tian H, Yue K, Liu J, Zhang B, Li X, Ding Z (2016) A P-Loop NTPase regulates quiescent center cell division and distal stem cell identity through the regulation of ROS homeostasis in *Arabidopsis* root. PLoS Genet 12: e1006175

Zhu YL, Pilon-Smits EA, Jouanin L, Terry N (1999) Overexpression of glutathione synthetase in indian mustard enhances cadmium accumulation and tolerance. Plant Physiol 119: 73–80

Zolotarev AS, Unnikrishnan M, Shmukler BE, Clark JS, Vandorpe DH, Grigorieff N, Rubin EJ, Alper SL (2008) Increased sulfate uptake by *E. coli* overexpressing the SLC26-related SulP protein Rv1739c from *Mycobacterium tuberculosis*. Comp Biochem Physiol A Mol Integr Physiol 149: 255–266

